# The trait coding rule in phenotype space

**DOI:** 10.1101/2022.09.29.510032

**Authors:** Jianguo Wang, Xionglei He

## Abstract

Genotype and phenotype are both the themes of modern biology. Despite the elegant protein coding rules recognized decades ago in genotype, little is known on how traits are coded in a phenotype space (*P*). Mathematically, *P* can be partitioned into a subspace determined by genetic factors (*P*^G^) and a subspace affected by non-genetic factors (*P*^NG^). Evolutionary theory predicts *P*^G^ is composed of limited dimensions while *P*^NG^ may have infinite dimensions, which suggests a dimension decomposition method, termed as uncorrelation-based high-dimensional dependence (UBHDD), to separate them. We applied UBHDD to a yeast phenotype space comprising ~400 traits in ~1,000 individuals. The obtained tentative *P*^G^ matches the actual genetic components of the yeast traits, explains the broad-sense heritability, and facilitates the mapping of quantitative trait loci, suggesting the tentative *P*^G^ be the yeast genetic subspace. A limited number of latent dimensions in the *P*^G^ were found to be recurrently used for coding the diverse yeast traits, while dimensions in the *P*^NG^ tend to be trait specific and increase constantly with trait sampling. A similar separation success was achieved when applying UBHDD to the UK Biobank human brain phenotype space that comprises ~700 traits in ~26,000 individuals. The obtained *P*^G^ helped elucidate the genetic versus non-genetic origins of the left-right asymmetry of human brain, and reveal several hundred novel genetic correlations between brain regions and dozens of mental traits/diseases. In sum, by developing a dimension decomposition method we show that phenotypic traits are coded by a limited number of genetically determined common dimensions and unlimited trait-specific dimensions shaped by non-genetic factors, a rule fundamental to the emerging field of phenomics.

## Introduction

The physical world is both macroscopic and microscopic, the former of which is the manifestation of the latter. Physicists adopt two rather parallel frameworks to describe the world: classical mechanics for the macroscopic layer and quantum mechanics for the microscopic layer^1^. For biologists, the macroscopic layer is phenotype and the microscopic layer is genotype. The mainstream of current biology adopts a bottom-up thinking: because genotype is the basis of phenotype, we rely on the former to understand the latter^2^. However, efforts of applying genotype to understanding phenotype appear successful only for rather simple phenotypic traits^3–5^. Hence, a possible complement to biologists is, like what the physicists used to do, to discover the rules working at the macroscopic layer (i.e, phenotype)^6,7^. As a matter of fact, many interesting patterns regarding the dimension sharing, coordination, and trade-off among phenotypic traits have been discovered in various organisms^8–14^. By focusing on specific traits and specific organisms these discoveries are, however, far from sufficient for constituting a satisfactory framework for understanding phenotype. The recent availability of large-scale phenomic data in a variety of species^15–18^ motivated us to seek for more general rules working at the phenotypic layer.

Phenotype is affected by both genetic and non-genetic (including environmental) factors. In quantitative genetics a phenotypic trait can be mathematically partitioned as^19^:

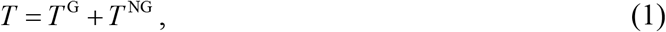

where *T* represents a focal trait, *T*^G^ is the genetic component fully determined by genotype, and *T*^NG^ is the residual component likely affected by environmental variables, developmental plasticity, measuring errors, human definitions (Supplementary Note I), and so on, collectively termed as non-<u>g</u>enetic factors. *T*^G^ contributes to the broad-sense heritability 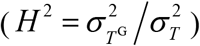 of *T*, and can be estimated by mathematical methods such as linear mixed model (LMM) when biological replicates are available^19^. *T*, *T*^G^ and *T*^NG^ are all vectors if a population is examined. When all phenotypic traits of a species are considered, we have:

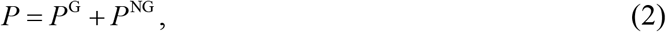

where *P* represents the phenotype space formed by all *T*, *P*^G^ represents the genetic subspace formed by all *T*^G^, and *P*^NG^ represents the residual (or non-genetic) subspace formed by all *T*^NG^. Specifically, *P*, *P*^G^ and *P*^NG^ are each a multi-dimensional linear space described by a matrix in which columns are trait vectors. Following the matrix notation there exists a set of orthogonal base vectors in *P*^G^, which we term as G-dimensions. Linear combinations of the G-dimensions can form all vectors in *P*^G^ (i.e., all *T*^G^). Similarly, the NG-dimensions in *P*^NG^ can be defined. Importantly, the number of G− (or NG−) dimensions is larger than or equal to the rank of *P*^G^ (or *P*^NG^). Accordingly, each trait *T* can be formulated as a linear function of the G-dimensions and NG-dimensions:

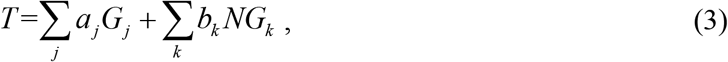

where *G_j_* represents the j^th^ G-dimension in *P*^G^, *NG_k_* represents the k^th^ NG-dimension in *P*^NG^, and *a_j_* and *b_k_* represent the coefficients of *G_j_* and *NG_k_*, respectively. Apparently, 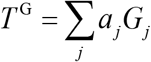 and 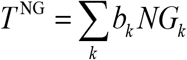. To be clear, throughout the paper the genetic component and non-genetic component of a trait *T* refer strictly to *T*^G^ and *T*^NG^, respectively.

The Fisher’s geometric model of evolutionary adaptation^20^, together with the extension by Orr^21^ and others^22,23^, predicts that the number of G-dimensions in *P*^G^ should be rather small for extant organisms. This is because a very large number of G-dimensions would hinder the adaptation to new environments, leading to extinction of the organisms, a phenomenon termed as ‘cost of complexity’^21^. Although the model does not predict exactly how small the number of G-dimensions should be^24^, we are still strongly inspired to hypothesize a limited number of G-dimensions^25^. In sharp contrast, the number of NG-dimensions in *P*^NG^ would be infinite. This is because of the variability of environment, the randomness of developmental plasticity and measuring error, and the arbitrariness of human definition^7^. The enormous complexity resulting from the infinite dimensionality of *P*^NG^ suggests the necessity of separating *P*^G^ from *P*^NG^ before revealing any rules in *P*.

In this study we started with asking how marginal correlation represents high-dimensional dependence in a multi-dimensional space. The answer enabled us to design a geometric method for separating two subspaces with distinct dimensionality. The method offered a phenome-based approach to separating a yeast phenotype space and a human brain phenotype space, respectively. The separated tentative genetic and non-genetic subspaces were then validated by available experimental benchmarks. The separation results revealed a rather simple geometric rule on how traits are coded in phenotype space. The results also provided novel phenotypic understandings not only within human brain but between brain regions and a variety of mental traits/diseases. In addition, this study developed a novel dimension decomposition strategy for dealing with the “curse of dimensionality”.

## Results

### Theory of uncorrelation-based high-dimensional dependence (UBHDD)

Let’s first consider a two-dimensional space with three non-parallel and non-orthogonal vectors *α*, *β*, and *η* (Fig. 1). Based on linear algebra, *η* can always be expressed as a linear function of *α* and *β* no matter whether *α* and *β* have strong (Fig. 1a) or little (Fig. 1b) marginal correlation (for simplicity, correlation, measured by Pearson’s correlation coefficient throughout the paper) with *η*. This is because the three vectors share the two dimensions (X-axis and Y-axis). In the three-dimensional space shown in Fig. 1c, *η* has a unique dimension (Z-axis). As a result, *η* can no longer be expressed by *α* and *β* despite the same correlations with *α* and *β* as in Fig. 1b. Hence, dimension sharing but not correlation underlies the high-dimensional dependence among vectors.

**Fig. 1.**
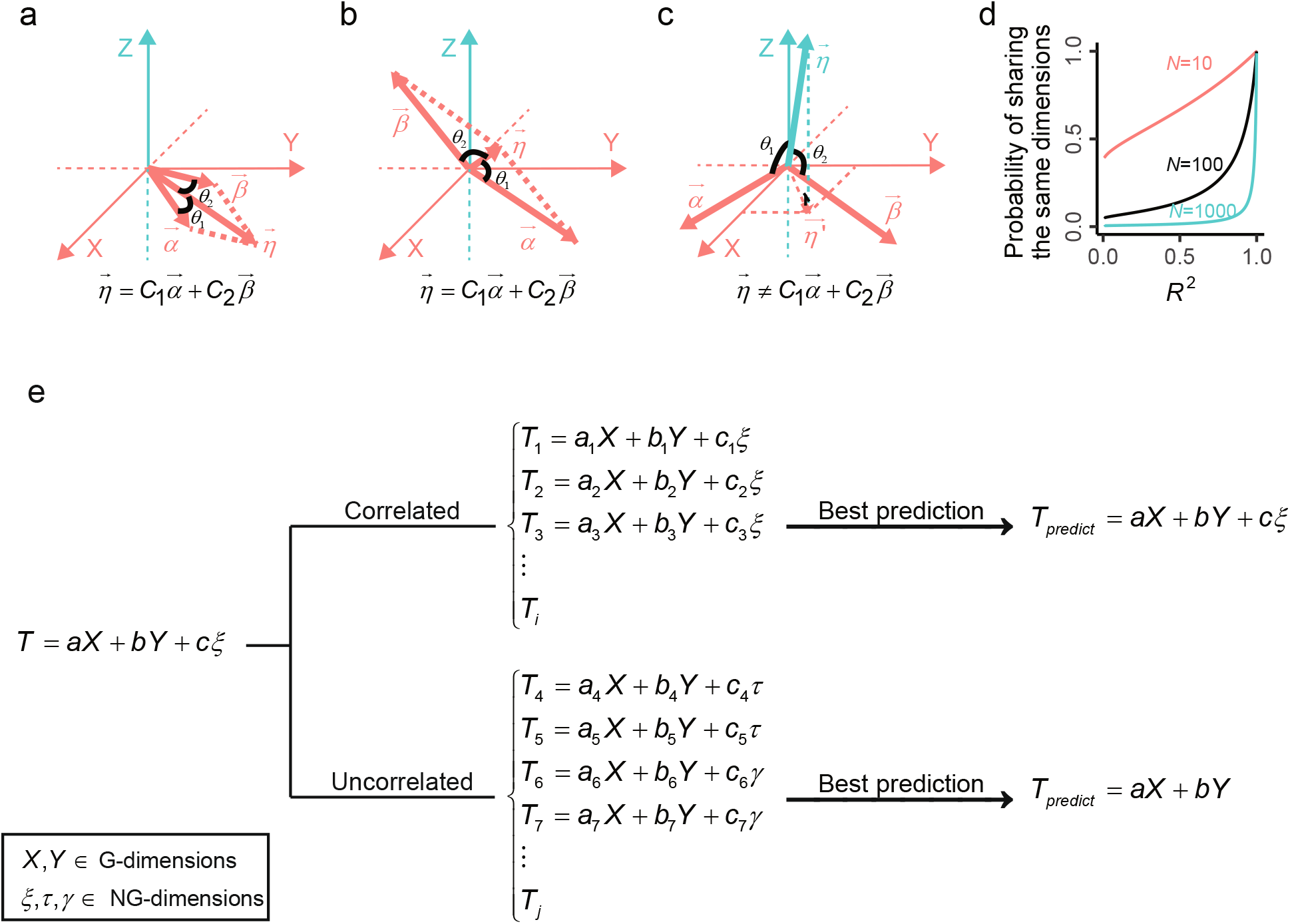
The underlying theory of UBHDD. **(a,b):** In each panel there are three non-parallel and non-perpendicular vectors (*α*, *β*, *η*) shown in a two-dimensional plane defined by the X and Y axes. Vector *η* can always be expressed as a linear combination of vectors *α* and *β* no matter whether the angle *θ*_1_ (*θ*_2_) between *η* and *α* (*β*) is small (i.e., correlated) as in panel a, or close to 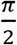 (i.e., uncorrelated) as in panel b, where 0 < 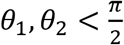. Note that, with two traits represented by two vectors, the correlation (Pearson’s *R*) between the two traits is equal to the cosine of the angle (*θ*) between the two vectors (i.e., *R* = cos*θ*). **(c):** Three non-parallel and non-perpendicular vectors in a three-dimensional space are shown, in which *η* is nearly perpendicular to the XY-plane and thus uncorrelated to *α* and *β*. In contrast to panel b, here *η* can no longer be expressed as a linear combination of *α* and *β* because *η* owns a unique dimension (Z-axis). **(d):** Uncorrelated vectors have a moderate probability of sharing the same dimensions in a space of low dimensionality. The probability of sharing dimensions depends the dimensionality of space (*N*), the dimensionality of each trait (*k*), and the correlation level (Pearson’s *R*^2^, 0<*R*^2^<1) of the two vectors. Based on a general geometric deduction (Supplementary Note II), the probability would converge to one when *R*^2^ converges to one, and to zero when *N* converges to infinity. Here the probability trajectories as a function of *R*^2^ are shown for *N*=10, 100 and 1000, respectively, with *k*=2 for both vectors. For a very small *R*^2^ the probability approaches zero for *N*=1000 but remains a moderate level for *N*=10. **(e):** A simple example shows how UBHDD would work. A trait *T* is formed by two dimensions of *P*^G^ (*X* and *Y*) and one NG-dimension of *P*^NG^ (*ξ*), where the dimensionality *N* is small for *P*^G^ but very large for *P*^NG^. Following the theoretical deduction in panel **d**, the correlated traits of *T* would have the same G- and NG-dimensions as *T*. Hence, the best model of predicting *T* using its correlated traits would still be a linear function of *X*, *Y* and *ξ*. In other words, the G- and NG-dimensions are not separable using correlated traits. In contrast, the uncorrelated traits of *T* would likely share G-dimensions but not NG-dimensions with *T* because of the dimensionality disparity between *P*^G^ and *P*^NG^. As a consequence, the best model of predicting *T* using its uncorrelated traits would represent only the genetic component of *T*.

We derived the probability (*Pr*(Ψ)) of two *k*-dimensional vectors that share the same dimensions in an *N*-dimensional space as a function of their correlation (Supplementary Note II). Without loss of generality, the probability trajectories of *N* = 10, 100, and 1,000 are shown, respectively, for two vectors with *k* = 2 (Fig. 1d). There are three corollaries: First, the probability converges to one if the two vectors have a strong correlation for any finite *N*, which is formulated as *Pr*(Ψ) → 1 if *R*^2^ → 1 and *N* < *N*_0_, where *N*_0_ is a finite number. Second, with the decrease of the correlation between the two vectors, the probability converges to zero in a space of very large *N*, which is formulated as *Pr*(Ψ) → 0 if *N* → ∞ and 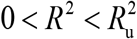, where 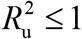. Third, with the decrease of the correlation between the two vectors, the probability remains reasonably high in a space of small *N* (e.g., *N* = 10), which is formulated as *Pr*(Ψ) > *Pr*_0_ if *N* < *N*_0_ and 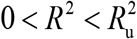, where *Pr*_0_ ≥ 0 . Accordingly, given an *R*_u_ with a small absolute value, uncorrelated vectors 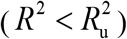 would have a rather high probability of sharing dimensions in a space of small *N* but little probability in a space of very large *N*. This suggests a strategy for separating *P*^G^ from *P*^NG^, the former of which is hypothesized to have a limited *N* while the latter an infinitely large *N*.

Fig. 1e shows how to model a trait *T* that is a function of G-dimensions and NG-dimensions in a given *P*. Because its correlated traits likely have the same G- and NG-dimensions as *T*, the best model of predicting *T* by its correlated traits would approximate the whole *T*. This way, the genetic and non-genetic components of *T* cannot be separated. In contrast, the uncorrelated traits of *T* would likely share G-dimensions but not NG-dimensions with *T* according to the deduction in Fig. 1d, if the dimensionality *N* is much smaller in *P*^G^ than in *P*^NG^. As a result, the best model of predicting *T* by its uncorrelated traits would represent only the genetic component of *T*. The residue (*T* − *T*_predict_) would then be the non-genetic component *T*^NG^. Because *P* is a collection of traits, by conducting such uncorrelation-based separation for every trait in *P* we would achieve the separation of *P*^G^ from *P*^NG^. We term the method <u>u</u>ncorrelation-<u>b</u>ased <u>h</u>igh-<u>d</u>imensional <u>d</u>ependence (UBHDD).

### Validation of UBHDD using simulation

To test if UBHDD can separate subspaces of distinct dimensionality, we simulated a space *P* that comprises a subspace *P*^G^ with a small number of G-dimensions (*N*_1_=10) and another subspace *P*^NG^ with a much larger number of NG-dimensions (*N*_2_=10,000) (Supplementary Note III). The G-dimensions and NG-dimensions are generated by standard multivariate normal distribution. Each trait (*T*) is generated by random linear combination of the G-dimensions and NG-dimensions as given by Eq. (3), with the former representing *T*^G^ and the latter representing *T*^NG^. A total of 1,000 traits are simulated in a population of 1,000 individuals. Each trait is standardized such that the variance of *T*^G^ equals to the broad-sense heritability (*H*^2^). Combining all *T*^G^ or all *T*^NG^ forms the sampled *P*^G^ or *P*^NG^, respectively.

UBHDD is conducted as follows (Methods): For all possible trait pairs two traits are defined as uncorrelated if their Pearson’s 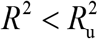, where 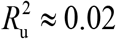, corresponding to *p* = 0.01 (t-test with Bonferroni correction); with conventional machine learning framework (LASSO) we modeled a trait *T* using its all uncorrelated traits; the predicted vector and the residual vector, designated as *T*^g^ and *T*^ng^, approximate the genetic component *T*^G^ and non-genetic component *T*^NG^, respectively; the resulting matrices containing all *T*^g^ or all *T*^ng^ are called *P*^g^ or *P*^ng^, approximating *P*^G^ and *P*^NG^, respectively.

As expected, with the increase of trait sampling the number of sampled dimensions is much more rapidly saturated for *P*^G^ than *P*^NG^ (Fig. 2a). We noted that the sampled dimensions in *P*^NG^ would keep increasing if the dimensionality of *P*^NG^ were infinitely large. Two correlated traits often share both G-dimensions and NG-dimensions while two uncorrelated traits could share G-dimensions but rarely NG-dimensions (Fig. 2b). This suggests G-dimensions but not NG-dimensions would underlie the signal of UBHDD. Indeed, in all cases we found the *T*^g^ obtained by UBHDD highly correlated with *T*^G^, the actual genetic component of *T* (Fig. 2c). The variance of *T*^g^ also matches well the variance of *T*^G^, the broad-sense heritability of *T* (Fig. 2d). We also simulated spaces with *N*_1_ = 20, 50, or 100 (*N*_2_ remains unchanged), and obtained largely the same results (Fig. S1). These analyses validated the capacity of UBHDD in separating *P*^G^ from *P*^NG^.

**Fig. 2.**
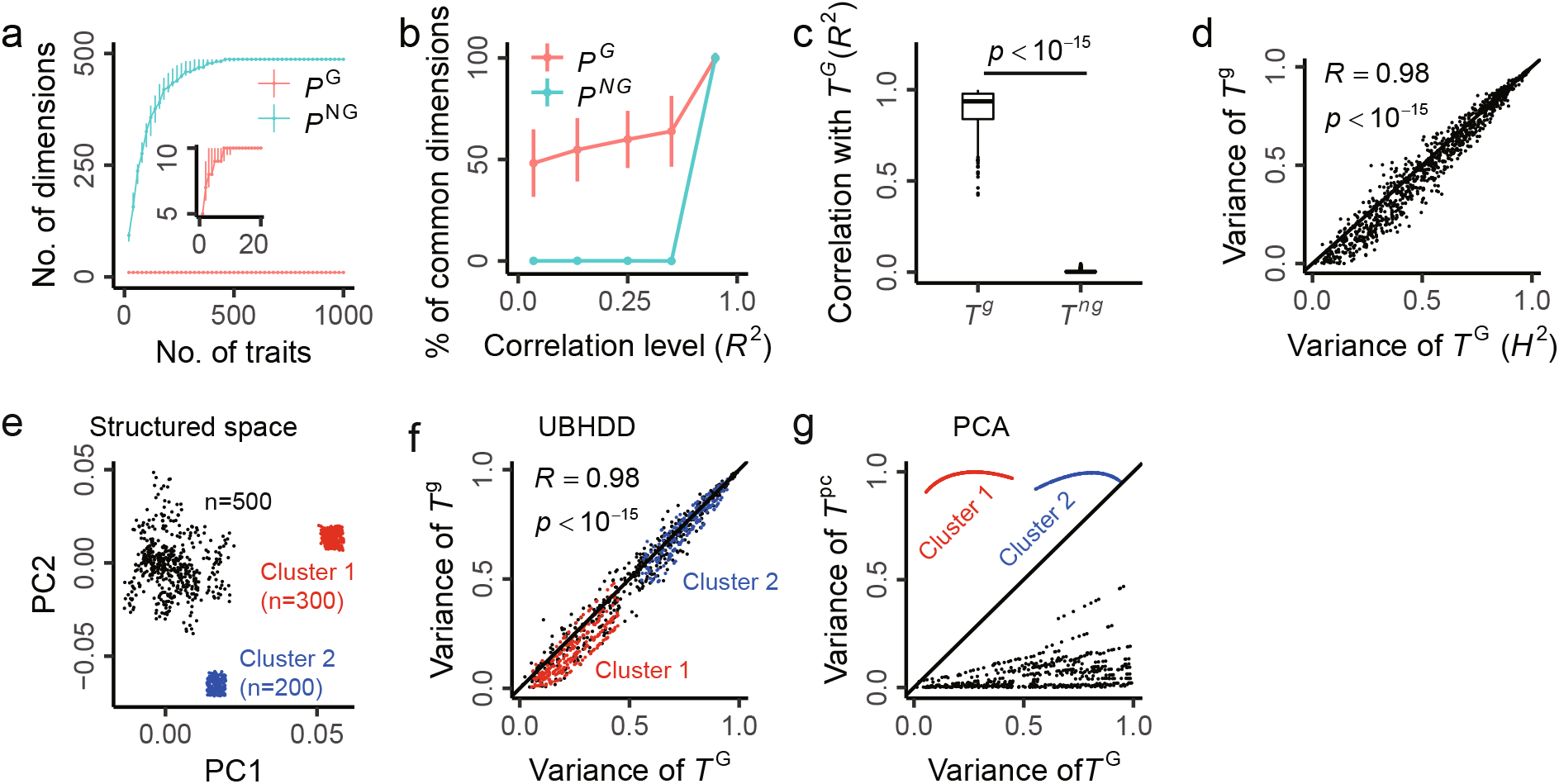
Separation of *T*g from *T*ng by UBHDD in simulated phenotype spaces. **(a):** In the simulated phenotype space the number of obtained dimensions is saturated much more rapidly in *P*^G^ (*N*_1_=10) than in *P*^NG^ (*N*_2_=10,000) as a function of trait sampling. Error bar represents the 95% confidence interval estimated by 100 replicates of trait sampling. **(b):** The probability of sharing *P*^G^ dimensions remains constantly high for traits with various levels of correlation. In contrast, the probability of sharing *P*^NG^ dimensions rapidly converges to zero for traits of low correlation. The threshold *R*_u_~= 0.15 (0.147) is used for defining uncorrelated traits in the simulated phenotype space, which corresponds to *p* = 0.01 after correction for multiple testing. Error bar shows the standard error. **(c):** The *T*^g^ obtained by UBHDD is highly correlated with the actual genetic component *T*^G^ of the simulated traits. As a control, the correlation between *T*^ng^ and *T*^G^ is also shown. A total of 1,000 traits are examined and standard box plots are used to display the data. The *p*-value is computed by paired t-test. **(d):** The variance of *T*^g^ in the simulated population is similar to the variance of *T*^G^, the actual broad-sense heritability (*H*^2^) of a trait in the population. Each dot represents a simulated trait and a total of 1,000 traits are shown. **(e):** The structure of a simulated structured phenotype space composed of 1,000 traits, with two large clusters comprising 300 and 200 highly correlated traits, respectively. Each dot represents a trait. **(f):** UBHDD has robust performance in the structured phenotype space, evidenced by the high similarity between the variance of *T*^g^ and the variance of *T*^G^. Each dot represents a trait and a total of 1,000 traits are shown. **(g)**: PCA is unable to reveal the genetic component of traits in the structured phenotype space. The top 2 PCs of the 1000 traits are used to model each trait (*T*^pc^), with the total explained variance comparable to that of *T*^G^. However, *T*^pc^ overfits the traits of the two large clusters and underfits the other traits.

It is worth noting that UBHDD is a method of dimension decomposition but not dimension reduction. We compared UBHDD with PCA, a classical dimension reduction method, in a simulated *P* with structure. The structured *P* was simulated as above except that two large clusters with strongly correlated members exist (Fig. 2e; Supplementary Note III). UBHDD remains successful in separating *P*^G^ from *P*^NG^, insensitive to the space structure (Fig. 2f). However, PCA overfits the traits in the two large clusters and underfits the others (Fig. 2g; Methods). The failure of PCA in separating *P*^G^ from *P*^NG^ is not surprising because PCA maximizes the explained variance of the top PCs and is therefore sensitive to data structure.

### Using UBHDD to separate a yeast phenotype space

We examined a phenotype space comprising 405 morphological traits of the budding yeast *Saccharomyces cerevisiae*^18^. The traits are measured in a population of 815 segregants, each of which has two clones/replicates and known genotype^26^ (Fig. 3a). The traits are typically about area, distance, angle, and brightness that describe the shape of mother cell and bud, the neck separating mother cell from bud, the localization of the nuclei in mother cell and bud, and so on, across different cell stages (Fig. 3b). The narrow-sense heritability (*h*^2^) of the traits ranges from 0 to 0.56 with a median of 0.15, and the broad-sense heritability (*H*^2^) ranges from 0 to 0.86 with a median of 0.42 (Fig. S2).

**Fig. 3.**
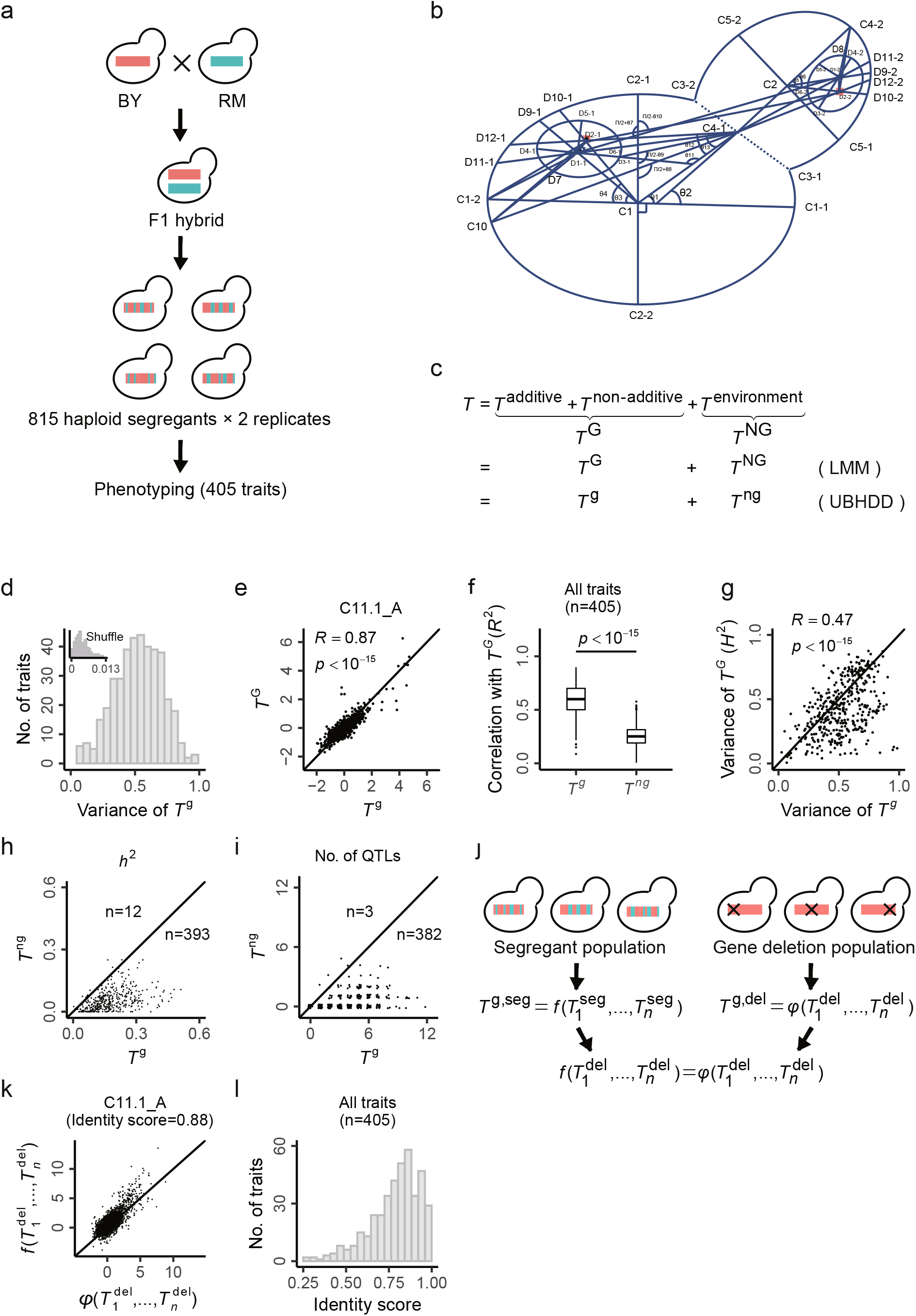
Separation of *T*^g^ from *T*^ng^ by UBHDD for 405 yeast traits. **(a):** A summary of the yeast phenome data. The yeast segregant population is generated by a cross of two *S. cerevisiae* strains (BY and RM). A total of 405 phenotypic traits are characterized for each of 815 segregants, with two clones examined for each segregant. **(b):** A schematic diagram of a yeast cell with landmarks for describing the shape or position of the cell wall and nuclei of the mother and daughter cells. **(c):** Two strategies used for separating the genetic component from the non-genetic component of a quantitative trait. The genotype-based linear mixed model (LMM) is a classical strategy, and the resulting components are denoted as *T*^G^ and *T*^NG^. The phenome-based UBHDD, which requires no genotype information, is proposed in this study; the resulting components are denoted as *T*^g^ and *T*^ng^. **(d):** Substantial trait variance is captured by *T*^g^. The inset shows the results of shuffling analyses that serve as a negative control for the UBHDD signals. Note that the total variance of a trait is one. **(e-f):** The *T*^g^ estimated by UBHDD is highly similar to the *T*^G^ derived from LMM. As a control, *T*^ng^ is distinct from *T*^G^. The panels e shows the details of a randomly selected trait (C11.1_A), and the panel f shows the summary for the 405 traits. **(g):** The variance of *T*^g^ is similar to the variance of *T*^G^, the broad-sense heritability of *T* estimated by LMM. **(h):** The narrow-sense heritability (*h*^2^) of *T*^g^ is generally larger than that of *T*^ng^. Each dot represents a trait and 405 yeast traits are examined. **(i):** There are more QTLs detected in *T*^g^ than *T*^ng^. Each dot represents a trait and 405 traits are examined (with 20 traits on the line of y=x). The dots are plotted with jitter for visual distinguishability. **(j):** For a given trait the *T*^g^ function learned in the segregant population can be compared with the *T*^g^ function learned in another yeast population comprising 4,718 gene deletion strains, which assesses the robustness of UBHDD. **(k):** For a randomly selected trait C11.1_A, the two functions produce highly similar *T*^g^ in the gene deletion strains, with an identity score = 0.88. The identity score is defined as the variance component along y=x in the scatter plot. Each dot represents a gene deletion strain and 4,718 strains are examined. **(l):** Density distribution of the identity scores of the 405 yeast traits.

Since biological replicates are available for the yeast phenome, we can use linear mixed model (LMM) to separate the *T*^G^ from *T*^NG^ for each of the traits. Meanwhile, the separation could be done by UBHDD, which requires only phenome information according to the above theory and simulation results (Fig. 3c). We will then use the results of LMM to benchmark UBHDD.

We applied UBHDD to the 405 yeast traits and obtain for each of them the *T*^g^ and *T*^ng^ (Methods). The obtained *T*^g^ explains trait variance at a level ranging from 0.03 to 0.98, with a median=0.53 among all traits (Fig. 3d). Hence, strong high-dimensional dependence between the uncorrelated yeast traits is observed. To assess the potential false positive/background signals, we conducted shuffling analyses by randomly swapping the focal trait values among individuals while maintaining the uncorrelated traits unchanged (Methods). We found virtually no trait variance explained (maximum=0.013 among all traits) by the *T*^g^ obtained in the shuffled dataset (Fig. 3d). Hence, technical biases in the UBHDD modeling process are negligible. Notably, the results of the shuffling analyses are actually consistent with our intuition in the empirical world that uncorrelated objects are independent, which has a hidden assumption for infinite dimensionality. The observed strong UBHDD signals suggest a special set of latent dimensions underlying the yeast traits.

To test if the UBHDD signals represent actual genetic components, we applied LMM to separate *T*^G^ from *T*^NG^ for each trait by taking advantage of the replicate information (Methods). For most of the traits the UBHDD signal *T*^g^ is highly correlated to the actual genetic component *T*^G^ (Fig. 3e-f). The variance of *T*^g^ is comparable to the variance of *T*^G^, the broad-sense heritability estimated by LMM (Fig. 3g). The results are robust against the *R*u thresholds used for defining uncorrelated traits (Fig. S3). As another critical test, we expect *T*^g^ should have a larger narrow-sense heritability (*h*^2^) than *T*^ng^. Indeed, in most case the *h*^2^ of *T*^g^ is larger than that of *T*^ng^, and also more QTLs were detected for *T*^g^ than *T*^ng^ (Fig. 3h-i; Methods). Nevertheless, *T*^g^ is not identical to *T*^G^. The *T*^g^ estimation could be improved in a larger population that enables more robust UBHDD modelling; meanwhile, the *T*^G^ estimation could be more accurate if there were more than two replicates. Taken together, these results suggest the *T*^g^ obtained by UBHDD represents well the actual genetic components of the yeast traits.

### The separations by UBHDD are robust between two yeast populations

In addition to the segregant population (seg-population), we also examined a yeast gene-deletion population (del-population) that contains ~5,000 *S. cerevisiae* strains each lacking a non-essential gene (Fig. 3j). The same 405 traits are measured for each of the strains in the del-population^27^. We conducted UBHDD in the del-population and obtained the *T*^g^ and *T*^ng^ for each of the traits (Methods). We then compared the *T*^g^ functions learned in del-population with the *T*^g^ functions previously learned in seg-population (Methods). Taking the trait C11.1_A as an example, when the *T*^g^ function learned in seg-population is applied to del-population, the *T*^g^ estimations are highly similar to the estimations by the *T*^g^ function learned in del-population, with an identity score = 0.88 (Fig. 3k; Methods). The identity score of the 405 traits ranges from 0.29 to 0.99 with a median=0.82 (Fig. 3l), suggesting the genetic subspace obtained by UBHDD be robust between the two yeast populations.

### Using UBHDD to separate human brain phenotype space

To test if UBHDD works in a more complex phenotype space, we examined UK Biobank human phenome. We focused on the 675 image-derived phenotypes (IDPs) of brain generated by dMRI in 25,957 white British individuals without kinship and with genotype available (Fig. 4a; Methods)^28^. These brain image traits represent nine different measures including fractional anisotropy (FA), intra-cellular volume fraction (ICVF), isotropic or free water volume fraction (ISOVF), mean diffusivity (MD), diffusion tensor mode (MO), orientation dispersion index (OD) and the three eigenvalues in a diffusion tensor fit (L1, L2 and L3) in up to 75 brain regions.

**Fig. 4.**
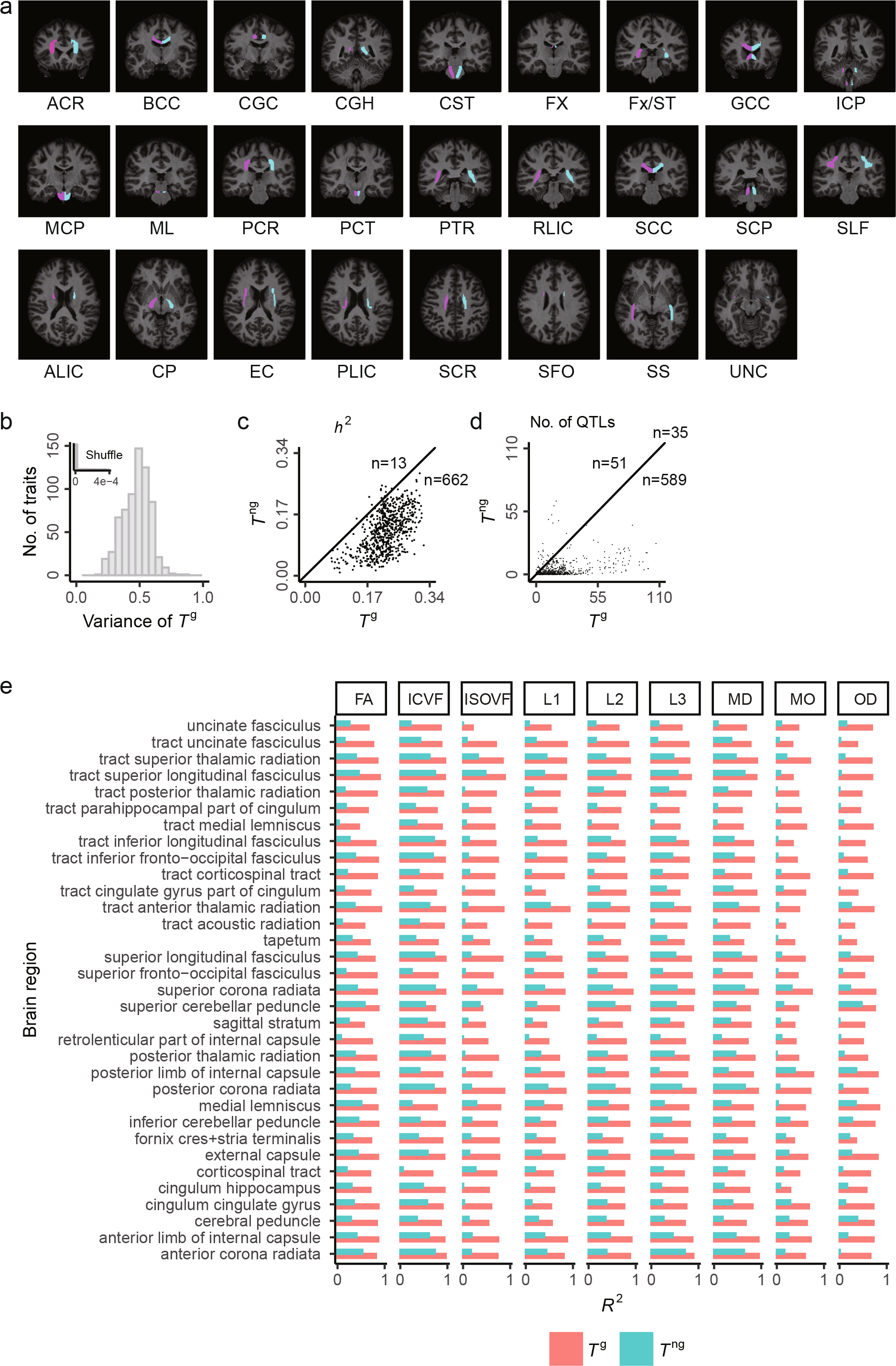
Separation of *T*g from *T*ng by UBHDD for 675 human brain image traits. **(a):** A summary of the human brain phenome data. Here the typical brain regions in the classical brain region atlas of Johns Hopkins (used in UK biobank) are shown. For example, ACR is short for anterior corona radiate and others are listed in Methods. **(b):** Substantial trait variance is captured by *T*^g^. The inset shows the results after randomly shuffling the individuals. Note that the total variance of a trait is one. **(c):** The *h*^2^ of *T*^g^ is generally larger than that of *T*^ng^. Each dot represents a trait. **(d):** There are more QTLs detected in *T*^g^ than *T*^ng^. Each dot represents a trait. **(e):** The left-right similarity is invariably stronger in *T*^g^ than *T*^ng^, suggesting non-genetic factors be the major source of brain left-right asymmetry. The similarity is measured by Pearson’s *R*^2^ between a symmetrical trait pair. A total of 297 trait pairs are examined.

We applied UBHDD to the 675 brain image traits after excluding covariates and obtained for each of them the *T*^g^ and *T*^ng^ (Methods). The obtained *T*^g^ explains trait variance at a level ranging from 0.17 to 0.87, with a median=0.48 among all traits (Fig. 4b). We conducted the same shuffling analysis as in yeast and again found virtually no trait variance (maximum=4e-4 among all traits) explained by the *T*^g^ obtained in the shuffled dataset (Methods). The results are robust against the *R*_u_ thresholds for defining uncorrelated traits (Fig. S4). Because there are, unlike yeasts, no clones (i.e., monozygotic twins) for most individuals, we couldn’t use LMM to estimate *T*^G^ and broad-sense heritability. Instead, we examined narrow-sense heritability. Consistent with the findings in yeast, *T*^g^ in general has a larger *h*^2^ than *T*^ng^; there are also more QTLs detected in *T*^g^ than *T*^ng^ (Fig. 4c-d; Methods). Notably, for those traits with a strong enrichment of the additive variance in *T*^g^, the number of QTLs of *T*^g^ is even larger than that of the whole trait *T*, suggesting novel genetic basis revealed by focusing on *T*^g^ (Fig. S5). These data suggest the *T*^g^ obtained here be at least enriched with the genetic components of the brain image traits. The results have two immediate applications.

First, it is helpful for addressing a long-standing puzzle, namely, the relative contribution of genetic versus non-genetic factors to the left-right asymmetry of human brain^29,30^. We examined all 297 symmetrical trait pairs each representing the same measure in two symmetrical brain regions. For each trait pair we calculated the Pearson’s *R*^2^ of *T*^g^ and *T*^ng^, respectively, among the individuals. In all trait pairs the *R*^2^ of *T*^g^ is much larger than that of *T*^ng^ (Fig. 4e). This finding suggests non-genetic factors be the major source of the brain asymmetry, highlighting environmental effects on asymmetry associated brain physiology and dysfunction.

Second, because of the enrichment of genetic component *T*^g^ should be particularly useful for identifying genetic correlations of the brain image traits with other traits including diseases. Such genetic correlations can inform the specific brain regions associated with or responsible for diseases, which would be valuable for diagnosis and/or therapy. We calculated genetic correlations^31^ between the 675 brain image traits and a curated set of traits with required summary statistics^32^. These traits include 33 common mental traits (including diseases and non-diseases), 13 respiratory/circulatory diseases that are associated with autonomic nervous system, and 32 miscellaneous diseases that do not seem to be tightly linked with brain (Methods; Table S1). A large number of statistically significant genetic correlations (*p*<0.05 after Benjamini-Hochberg correction for multiple testing; Methods) were detected with two notable features (Fig. 5a-c): First, the mental traits and the respiratory/circulatory diseases in general have more genetic correlations with the brain image traits than the miscellaneous diseases. Second, *T*^g^ performed much better than *T* in revealing genetic correlations. The results in turn support the enrichment of *T*^g^ for genetic component.

**Fig. 5.**
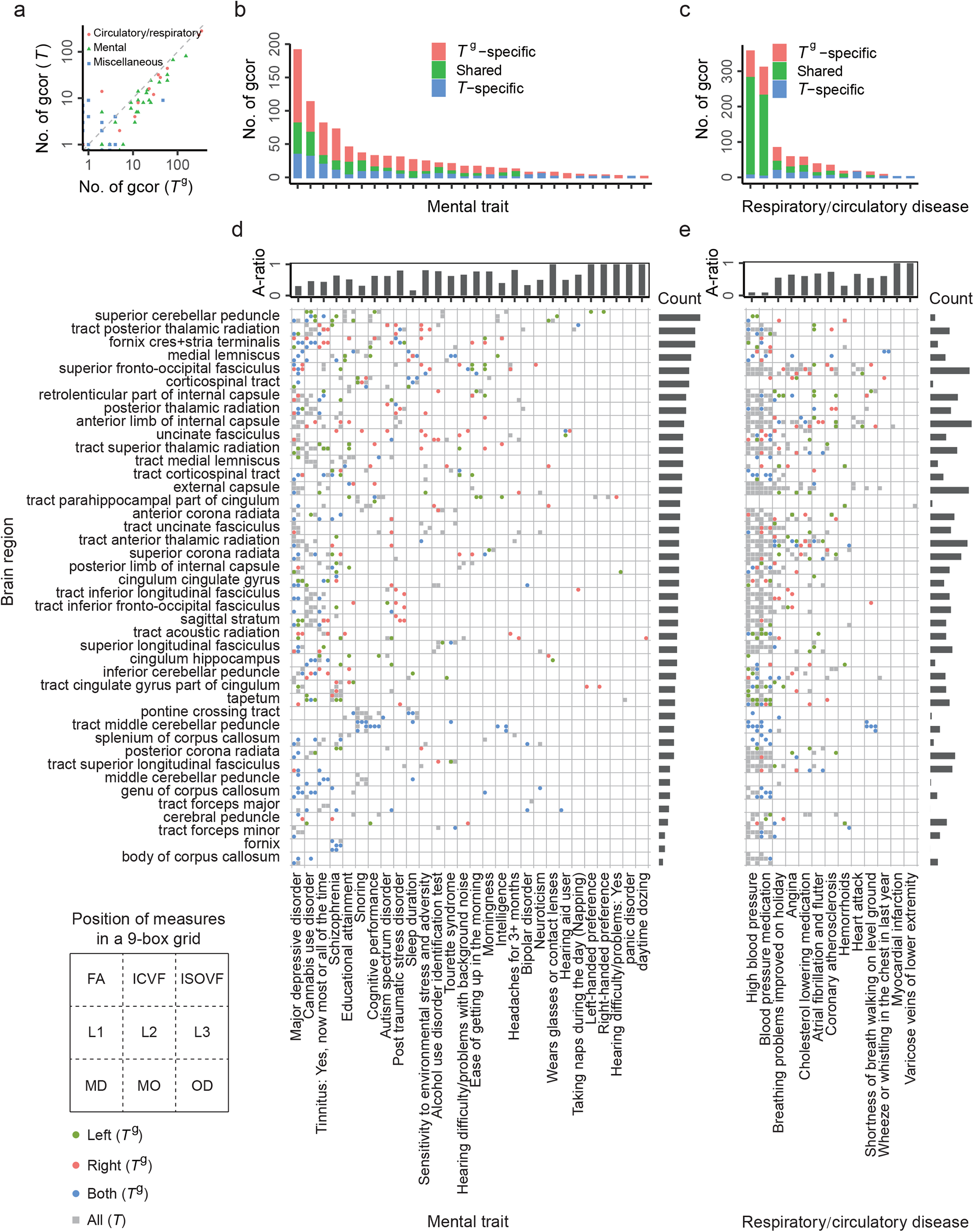
Novel genetic correlations revealed between brain image traits and mental traits/diseases. **(a):** The numbers of statistically significant genetic correlations with the 675 brain image traits (*T* versus *T*^g^) identified for each of mental traits/diseases. In general there are more genetic correlations identified with *T^g^* than with *T*. The ‘gcor’ is an abbreviation of genetic correlation. **(b):** The numbers of *T*^g^-specific, *T*-specific and shared genetic correlations for each of the mental traits. Five mental traits with no genetic correlations identified are excluded, leaving 28 that will be further examined. **(c):** Same as the panel b, except for respiratory/circulatory diseases. **(d):** All genetic correlations here identified between brain regions and the mental traits are shown. Each grid is a 9-box grid with each box representing a type of measure indicated at the bottom left corner. The colored dots show *T^g^*-specific genetic correlations, with the left/right hemisphere information provided. The grey square show all genetic correlations detected by *T*. The ‘A-ratio’ measures the number of genetic correlations with only one trait of a symmetrical trait pair divided by the total number of genetic correlations with all brain image traits. The ‘Count’ measures the total number of genetic correlations in a brain region relative to the sum of all brain regions. **(e):** Same as the panel d, except for respiratory/circulatory diseases.

To show more details we plotted all statistically significant genetic correlations for the mental traits and the respiratory/circulatory diseases, respectively (Fig. 5d-e). There are a few global patterns: First, brain regions vary substantially in the number and profile of correlated diseases/traits. For example, the brain region “fornix” has significant genetic correlation with only one disease Schizophrenia, while the region “superior fronto-occipital fasciculus” has significant genetic correlations with 19 diseases/traits. Second, diseases/traits vary substantially in the number and profile of correlated brain regions. For example, 13 out of 28 mental traits have significant genetic correlations with the brain region “tract parahippocampal part of cingulum”, while the number is 2 out of 13 respiratory/circulatory diseases. Third, the left and right brain hemispheres appear distinct for many diseases/traits. We computed an asymmetry ratio (A-ratio), which is the number of significant genetic correlations with only one trait of a symmetrical trait pair divided by the total number of significant genetic correlations detected, for each of the diseases/traits. There are many cases with a very large A-ratio, such as post-traumatic stress disorder and coronary atherosclerosis; meanwhile, there are salient cases with a very small A-ratio, such as sleep duration and high blood pressure. In addition to the global patterns, numerous specific understandings about the diseases can be updated. For example, previous studies reported statistically insignificant genetic correlations between the brain region “superior fronto-occipital fasciculus” and major depressive disorder (*p*~0.1)^33^, and between the brain region “superior cerebellar peduncle” and cannabis use disorder (*p*~0.2)^34^; in both cases we identified six image traits in the corresponding brain region showing significant genetic correlations with the corresponding disease. Five image traits in the brain region “tract middle cerebellar peduncle” are newly identified to have genetic correlations with the disease “shortness of breath walking on level ground”; interestingly, the same five traits are found to show genetic correlations with high blood pressure. For the traits educational attainment, cognitive performance and intelligence, there are nine, six and four newly identified brain regions, respectively. The rich novel information provided here would be of tremendous value for revealing the brain basis of the traits/diseases.

### Distinct dimensionality of *P*^g^ and *P*^ng^ reveals a trait coding rule

Using UBHDD we estimated the genetic component *T*^g^ and non-genetic component *T*^ng^ for each of the traits examined in yeasts and humans. Combining all *T*^g^ of the yeast traits (or human brain traits) forms *P*^g^, the estimated genetic subspace of the yeast (or human brain) phenotype space. Similarly, combining all *T*^ng^ forms *P*^ng^, the estimated non-genetic subspace. We then examined the latent dimensions in *P*^g^ and *P*^ng^, respectively. Using principal component analysis (PCA) we obtained the number of top PCs that explain 85% variance of a subspace. The cutoff (85% variance) was chosen because it approximated well the actual dimensionality of *P*^G^ in the simulated phenotype space analyzed in Fig. 2a-d (Fig. S6). We found that, with the increase of trait sampling, the number of PC dimensions is rapidly saturated for *P*^g^ but not for *P*^ng^ (Fig. 6a-b), highlighting the distinct dimensionality between *P*^g^ and *P*^ng^. The observed dimensionality disparity is consistent with the underlying theory of UBHDD.

**Fig. 6.**
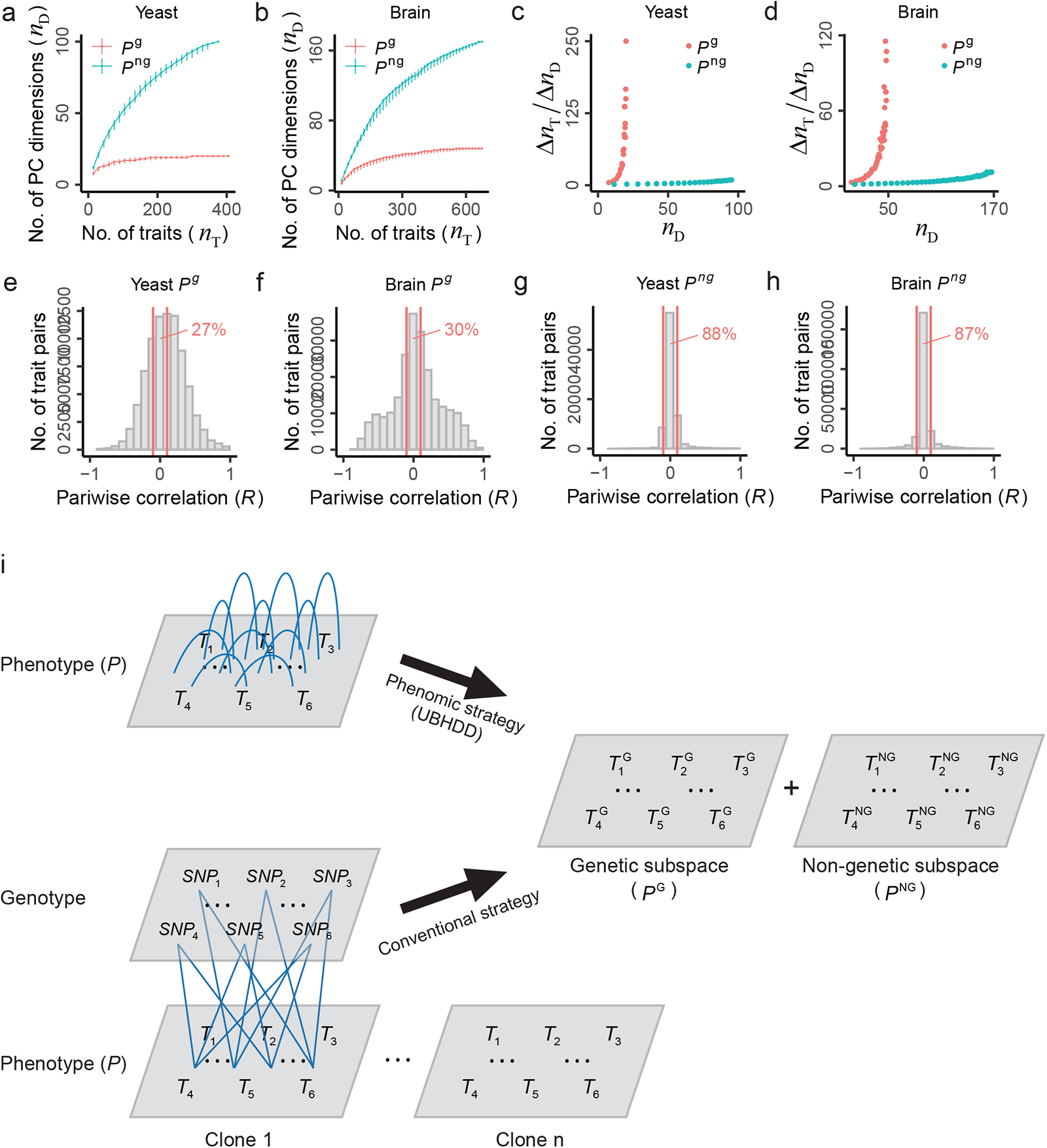
Distinct dimensionality of *P*^g^ and *P*^ng^. **(a-b):** The number of latent dimensions (*n*_D_) in *P*^g^ and *P*^ng^, respectively, as a function of the number of sampled traits (*n*_T_) for yeast (a) and human brain (b). The *n*D is estimated as the number of top principal components that explain 85% variance of the sampled traits. Error bar represents the 95% confidence interval estimated by 100 replicates of trait sampling. **(c-d):** The number of additional traits per additional dimension (Δ*n*_T_/Δ*n*_D_) is constantly high in *P*^g^ but small in *P*^ng^ for both yeast (c) and human brain (d). This suggests *P*^g^ dimensions be recurrently used to code the traits while *P*^ng^ dimensions tend to be trait-specific. **(e-f):** The genetic component (*T*^g^) of the traits often show correlation due to the common *P*^g^ dimensions. The Pearson’s *R* of *T*^g^ for all trait pairs is shown for yeast and human brain phenome, respectively. The two vertical red lines mark −0.1<*R*<0.1. **(g-h):** The non-genetic component (*T*^ng^) of the traits shows little correlation, echoing the fact that *P*^ng^ dimensions are trait-specific. The Pearson’s *R* of *T*^ng^ for all trait pairs is shown for yeast and human brain phenome, respectively. The two vertical red lines mark −0.1<*R*<0.1. **(i):** A phenome-based strategy (UBHDD) is proposed to decompose the genetic and non-genetic components of quantitative traits, which complements the kinship- or genotype-based conventional strategy.

To show how the dimensions of *P*^g^ and *P*^ng^ are used by the traits we calculated the gradient between the number of sampled traits (*n*_T_) and the number of obtained dimensions (*n*_D_), denoted as ∆*n*_T_/∆*n*_D_ . With the increase of dimensionality the gradient rapidly increases to be large for *P*^g^ but remains small for *P*^ng^, suggesting the *P*^g^ dimensions are recurrently used by the traits while the *P*^ng^ dimensions tend to be trait-specific (Fig. 6c-d). Consistently, the pairwise correlation of *T*^g^, which reflects dimension sharing between traits, is much larger than that of *T*^ng^ (Fig. 6e-h). Therefore, in both the yeast and human brain phenotype space the traits are coded by a rather small set of common dimensions that are determined by genotype and numerous trait-specific dimensions that are shaped by non-genetic factors.

## Discussion

Inspired by the evolutionary ‘cost of complexity’ theory in this study we designed a dimension decomposition method for separating subspaces of distinct dimensionality. We applied the method to a yeast phenotype space and a human brain phenotype space, respectively, to separate genetic subspace from non-genetic subspace. The separation results were then validated by available benchmarks. Despite the success, we cautioned that the results are just consistent with the evolutionary theory; resolving the debates on the theory^35,36^, which is beyond the scope of this study, requires further works.

The goal of this study is to find how traits are coded in phenotype space. Our analyses suggest phenotypic traits are coded by a limited number of genetically determined common dimensions and unlimited trait-specific dimensions that are shaped by non-genetic factors. The trait coding rule learned here underlies a phenome-based strategy for identifying the genetic component of a phenotypic trait (Fig. 6i). In addition to what we have presented, the strategy may help guide trait selection in future phenome mapping by gauging the captured genetic and non-genetic dimensions; it may also apply to the studies on the macroevolution of morphospace to extract the evolutionarily conserved genetic effects^12,14,37^.

There are a few technical issues worth discussing. First, the UBHDD method depends on dense sampling of a phenotype space. We may use a down-sampling strategy to assess the sufficiency of trait sampling. We found the overall performance of UBHDD for the yeast traits is nearly saturated (Fig. S7a); however, the performance for the human brain traits is sensitive to down-sampling (Fig. S7b), suggesting the current sampling of the brain space is still insufficient. Second, the uncorrelation thresholds (*R*_u_) used in this study may not be ideal. In principle, a smaller *R*_u_ is always helpful for avoiding the effects of non-genetic dimensions, which, however, would leave too few traits for conducting UBHDD. We found a good assessment of the threshold by examining the learned UBHDD functions. In our cases, the coefficients (Co) in the learned UBHDD function of a focal trait are not explained by the marginal correlations (Mc) of the explanatory variables (traits) to the focal trait (Fig. S8). In other words, the performance of UBHDD does not rely on those traits with stronger marginal correlation to the focal trait. In future, we may optimize *R*_u_ threshold trait by trait by considering the number of remaining traits under a given threshold as well as the relationship between Mc and Co. Third, only two phenotype spaces are examined. The generality of the findings should be further tested in more complex phenotype spaces. The last, but not the least, UBHDD offers a novel strategy for dealing with the “curse of dimensionality”^38,39^. Different from the conventional dimension reduction methods such as PCA, UBHDD works by assuming two types of latent dimensions in the space/system of interest. It is conceivable that, like phenotype space, many complex systems can be partitioned into a sub-system determined by intrinsic factors and another sub-system shaped by extrinsic factors, the former of which is of rather low dimensionality while the latter is composed of myriad dimensions. Hence, UBHDD could be a generally useful tool for studying a complex system.

## Methods

### Yeast segregant population (seg-population)

We study a panel of segregants of a yeast cross (*S. cerevisiae* strain BY × strain RM) generated by a previous study ^26^. A total of 1,008 segregants are available with genotypes, among which 815 were phenotyped ^18^. The obtained 405 phenotypic traits measure the areas and circumferences, the elliptical approximation, brightness, thickness, axis length, neck width, neck position, bud position, axis ratio, cell size ratio, outline ratio, proportion of budded cells, proportion of small budded cells, segment distances between mother tip, bud tip, middle point of neck, center of mother and bud, nuclear gravity centers, nuclear brightest points, angles between segments, and so on. The phenotyping was conducted for two clones of each segregant, and the trait values are Z-score transformed.

### Yeast single-gene-deletion population (del-population)

A previous study ^27^ has conducted similar phenotyping for 4718 yeast mutants each lacking a non-essential gene (del-population). The 405 traits in the seg-population are also available in the del-population. These traits in the del-population are scaled based on the mean and standard deviation in the seg-population to make models obtained from the two populations comparable.

### UK Biobank

We collect the 870 brain MRI phenotypes and related covariates (measuring center, age, sex, weight, home location and MRI system parameters) in UK Biobank^28^, of which 675 image-derived phenotypes (IDPs) measured by dMRI are chosen for UBHDD modelling, heritability estimation and QTL mapping. A few missing values in covariates are imputed by linear regression with brain phenotypes. A total of 25,957 White English without kinship and with genotypes are chosen. We conduct normalization on each of the 675 traits (R package, ‘bestNormalize’) and exclude the contribution of covariates by linear regression (R, ‘lm’). These traits are subject to following analysis including UBHDD, estimation of narrow-sense heritability and QTL mapping. We also obtain the genotypes for the subjects above. The pipeline is as follows. First, we use the software ‘qctool’ to extract SNPs (imputation score >0.8, MAF >0.01, genotype calling probability >0.9 and biallelic) for the 25,957 subjects above. Second, we use PLINK (beta 6.24, 6 Jun 2021) to extract SNPs (MAF >0.01, missing proportion of SNPs <0.1, Hardy-Weinberg Equilibrium exact test p-value >1e-6. After the two steps, we finally obtain the SNPs to be used to calculate narrow-sense heritability and conduct QTL mapping.

Typical brain regions: anterior corona radiata (ACR); anterior limb of internal capsule (ALIC); body of corpus callosum (BCC); cerebral peduncle (CP); cingulum cingulate gyrus (CGC); cingulum hippocampus (CGH); corticospinal tract (CST); external capsule (EC); fornix (FX); fornix cres+stria terminalis (Fx/ST); genu of corpus callosum (GCC); inferior cerebellar peduncle (ICP); medial lemniscus (ML); middle cerebellar peduncle (MCP); pontine crossing tract (PCT); posterior corona radiata (PCR); posterior limb of internal capsule (PLIC); posterior thalamic radiation (PTR); retrolenticular part of internal capsule (RLIC); sagittal stratum (SS); splenium of corpus callosum (SCC); superior cerebellar peduncle (SCP); superior corona radiata (SCR); superior fronto-occipital fasciculus (SFO); superior longitudinal fasciculus (SLF); uncinate fasciculus (UNC).

### Estimation of broad-sense heritability (*H*^2^) and *P*^G^ in yeast

A focal trait is modelled by linear mixed model (LMM) ^26^ as

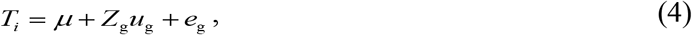

where *T_i_* is a focal trait, *μ* is the population mean, *Z*_g_ is the design matrix indicating which segregant each replicate belongs to, *u*_g_ is a vector of random effect, and *e*_g_ is a vector of residuals. The variance of the foal trait is decomposed into genetic effect 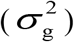 and environmental effect 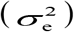. *H*^2^ is then estimated as

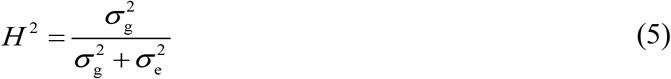

The random effect estimated for each segregant is defined as *P*^G^. The R package ‘lme4’ is used. Standard error is estimated by Jackknife.

### Estimation of narrow-sense heritability (*h*^2^) in yeast

A focal trait is modelled by LMM ^26^ as

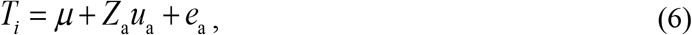

where *Z*_a_ is the identity matrix, *u*_a_ is a vector of random effect, 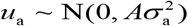, and *e*_a_ is a vector of residuals. The variance structure of the trait is formulated as

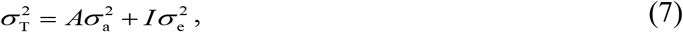

where *A* is estimated as the relatedness matrix, *I* is the identity matrix, 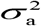 and 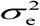 are additive variance and residual variance, respectively. Then, narrow-sense heritability (*h*^2^) is estimated as

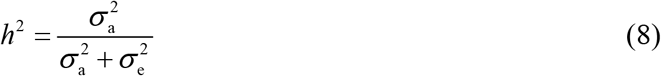

The R package ‘rrBLUP’ is used and standard error is estimated by Jackknife for yeast traits.

### QTL mapping in yeast

In yeast we follow the pipeline used in a previous study ^26^. The association between a focal trait and a focal SNP is calculated as LOD score defined by –n(ln(1-*R*^2^)/2ln(10)), where n is the number of non-missing segregants and *R* is the Pearson’s *R*. The threshold is determined by 1000 times shuffling of segregants. The strongest SNPs larger than the threshold in each chromosome are defined as QTLs. Total two rounds of QTL calling are conducted. The first round is conducted at the original traits and the second round is conducted at the residuals of the original traits. The R package ‘qtl’ is used.

### QTL mapping, heritability and genetic correlation in humans

In human, we use the widely used software GCTA^40^ to conduct QTL mapping. The threshold p is set to 5e-8. Clumping analysis is conducted by PLINK (beta 6.24, 6 Jun 2021) with the same p. The heritability of brain image traits is estimated by the R package ‘HDL’^41^. We collect the summary statistics of 78 traits/diseases with heritability larger than 0.01 estimated by ‘HDL’ from Center for Neurogenomics and Cognitive Research, Psychiatric Genomics Consortium, Social Science Genetic Association Consortium, UK Biobank and GWAS Catalog (Table S1). The genetic correlations between brain image traits and 78 diseases/traits are estimated by ‘HDL’ (Benjamini-Hochberg correction for multiple testing). Genetic correlations between mental traits are calculated by ‘HDL’.

### Uncorrelation-based high-dimensional dependence (UBHDD) modelling

The model is formulated as

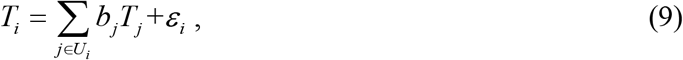

where *T_i_* is i^th^ trait, *b_j_* is the j^th^ coefficient, *ε*_*i*_ is the residual vector and *U_i_* contains the indices of uncorrelated traits of *Ti*. Then, we can obtain the estimated genetic component as

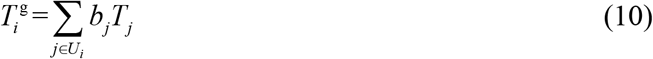

and the estimated non-genetic component as

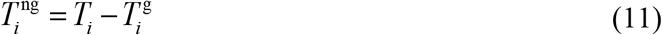

In yeast, the 815 samples are divided into a training subset with 715 samples and a testing subset with 100 samples. Then, a focal trait is modelled in the training subset. The prediction performance of the learned function was assessed as the *R*^2^ between predicted and observed trait values in the testing subset. Ten-fold cross validation and LASSO regularization are used to avoid overfitting (R package ‘glmnet’). Standard error is estimated by 20 repeats. In brain, the 25957 samples are randomly divided into 10 subsets with equal size, each time a subset is selected as a testing set and the others as a training set, and a focal trait is modelled in training set. The prediction performance of the learned function is assessed as *R*^2^ between predicted and observed trait values in the testing set. Ten-fold cross validation and LASSO regularization are used to avoid overfitting (Python package ‘glmnet’). Standard error is estimated by ten-fold cross validation. In simulated phenotype space and seg-population, the uncorrelation thresholds are set based on t-test with multiple-testing correction (*p*=0.01/(n-1) where n is the number of traits. In del-population, the threshold is set the same with that of seg-population to be comparable. In human brain, the threshold is set to be 0.15 by referring to the threshold in seg-population.

To control for potential technical bias, we also conduct shuffling analysis. For a focal trait *T_i_* in Eq. (9), we keep its uncorrelated traits *T_j_* unchanged and shuffle *T_i_* among individuals. Then, the same modelling process is conducted.

### Comparison between UBHDD and PCA in simulated structured population

We first simulated a structured population (Supplementary Note III). Then, we apply UBHDD to the structured population and obtain the *P*^g^. Next, we apply PCA to the structured population and keep the top PCs with explained variance up to the mean of those of UBHDD (the mean of variances of 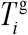). Finally, we recalculate the explained variance for each of simulated traits based on the kept top PCs.

### Identical Score

To evaluate the consistency of two variables, we can display them in a scatter plot. Then, the variance can be decomposed into two components, one along the straight line y=x and another along the straight line y=−x. This is similar to PCA except the transformed coordinate axes are pre-defined. The larger the variance of the component along y=x is, the more consistent the two variables are. The variance of the component along y=x is formulated as

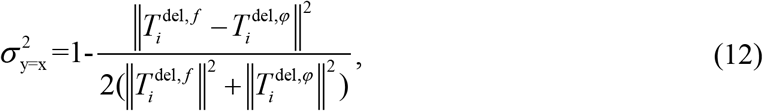

where 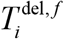 is the genetic component of a focal trait in del-population estimated by the function (*f_i_*) learned in seg-population; 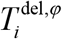 is the genetic component of a focal trait in del-population estimated by the function (*φ_i_*) learned in del-population. We name the 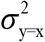 identity score to evaluate the robustness of *P*^g^ obtained by UBHDD between the two distinct yeast populations.

### Dimensionality estimation by PCA

For a subspace *P*^g^ or *P*^ng^, we conduct PCA after scaling (Z-score transformation). The number of top PCs with explained variance up to 85% is estimated as the dimensionality of the subspace.

## Code and data availability

The codes and the supporting data can be found at https://github.com/Jianguo-Wang/UBHDD.

## Supplementary Information

### Supplementary Note I

#### Non-genetic variation of complex traits can result from human definitions

Assuming a complex trait (*T*) is independently contributed by genetic factors (*G*) and a random noise (*E*), we have

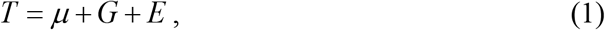

where *μ* ≠ 0 is the mean. Then, assuming the cube of *T* is also defined as a complex trait, we have

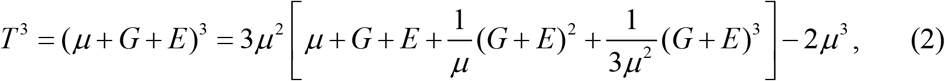

where new terms (*G* + *E*)^2^ and (*G* + *E*)^3^ are created by human definition. The new terms will contribute to the non-genetic variation.

Let us think the simplest situation, where *μ* = 5 (make sure *T* is positive), *G* is just a binary QTL encoded as {−1,1} with frequency equal to 0.5 and *E* follows standard normal distribution. A focal trait is defined by Eq. (1).

We conduct simulation and obtain the broad-sense heritability (*H*^2^) of *T* and *T*^3^ by linear mixed model (the same with that in Methods), shown as

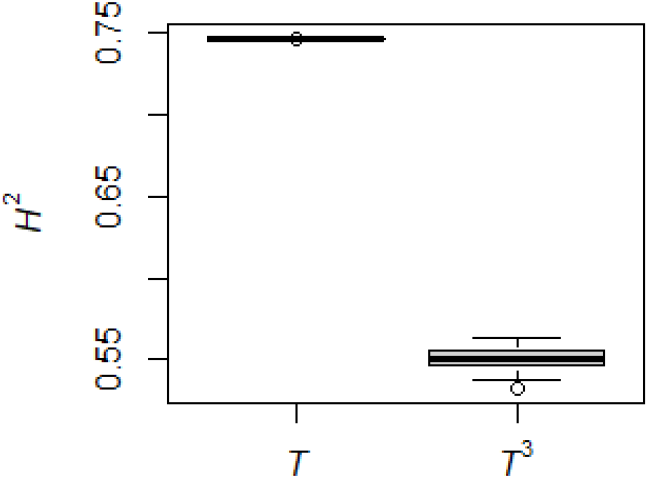

Therefore, human definitions can be a source of non-genetic variation.

### Supplementary Note II

#### The probability of two traits with the same dimensions

We achieve this estimation by transforming this problem into a geometric model of probability. We will first derive the general expression and then give the closed form in a specific condition. First, consider a general condition where two *k*-dimensional unit vectors *α* and *β* with angle equal to *θ* share *i* dimensions in an *N*-dimensional space (i.e., *α* and *β* each has *k* non-zero entries and *N-k* zero entries, sharing *i* non-zero entries), assuming 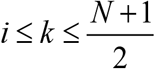 and 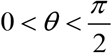 without loss of generality. In this condition, the first vector *α* has a form like 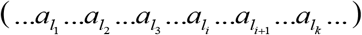 which has *k* terms not equal to zero and the second *β* has a form like 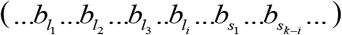 which also has *k* terms not equal to zero. Both of *α* and *β* have *i* terms not equal to zero in common *i* dimensions. Thus,

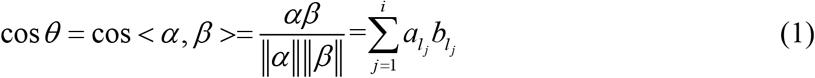

Let 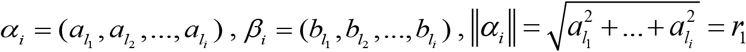 and 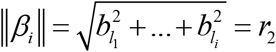.

Thus,

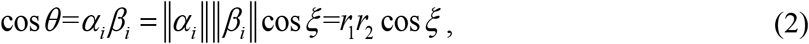

where 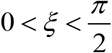 is the angle of *α_i_* and *β_i_*. Due to 0 < cos *θ* < 1, 0 < *r*_1_ ≤ 1, 0 < *r*_2_ ≤ 1, thus,

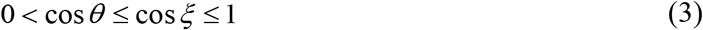

and

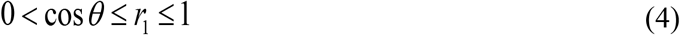

Set *r*_1_ = *r*, we obtain the geometric distribution of *α* as

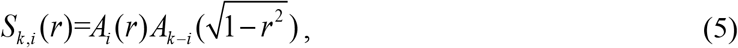

where *A_i_*(*r*) denotes the superficial area of an *i*-dimensional sphere with radius equal to *r* ^1^.

Because *α_i_* is the projected vector of *α* in the subspace of *β*, combining with Eq. (2), the projected vector *β*’ of *β* on *α_i_* satisfies

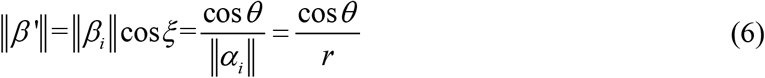

and

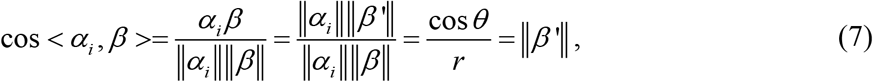

where *α_i_β* = ∥*α_i_*∥ ∥*β*’∥ and ∥*β*∥ = 1. Therefore, we obtain the geometric distribution of *β* as

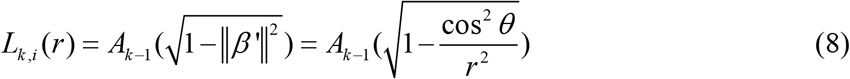

Therefore, combing Eq. (5) and Eq. (8), we obtain the geometric estimation for certain *α* and *β* given *k*, *i* and *θ* as

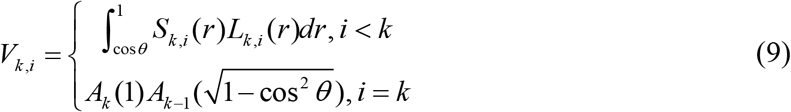

Therefore, the probability of two *k*-dimensional traits sharing the same dimensions (ψ) in an *N*-dimensional space given marginal correlation (cos*θ*) is estimated as

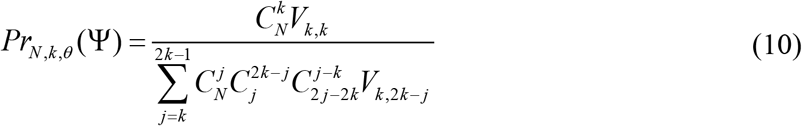

When *θ* → 0, there are *r* → 1 and *S_k,i_* (*r*) → 0 . Because 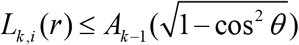, we get

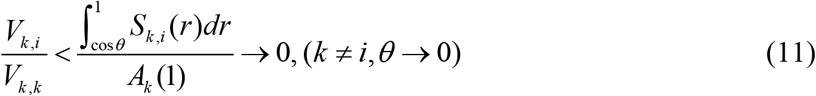

Therefore, given *N*

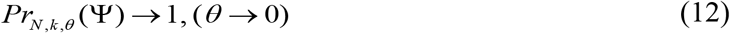

When *N* → ∞, there are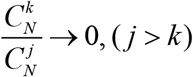, therefore, given *θ*

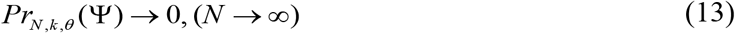

Next, consider a specific situation where two 2-dimensional vectors in an *N*-dimensional space. We can obtain

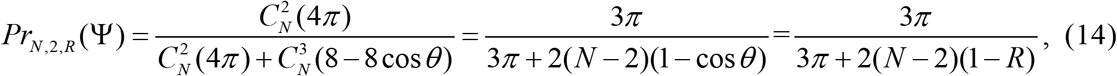

where *R*=cos*θ*. It is obvious that Eq. (14) still satisfies Eqs. (12–13). Eq. (14) is used to generate the trajectories in Fig. 1d. From Eq. (12–13) we can obtain three corollaries:

Corollary 1: *Pr*(Ψ) → 1 if *R*^2^ → 1 and *N* < *N*_0_, where *N*_0_ is a finite number.

Corollary 2: *Pr*(Ψ) → 0 if *N* → ∞ and 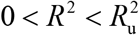, where 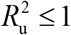.

Corollary 3: *Pr*(Ψ) > *Pr*_0_ if *N* < *N*_0_ and 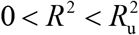, where *Pr*_0_ ≥ 0 .

Remarks. First, although *Pr_N,k,θ_*(Ψ) represents the probability of sharing the same dimensions, it’s hard to achieve linear deduction when few dimensions are shared in a very large space among a limited trait sample. Second, the corollary 1 guarantees that linear combinations of G-dimensions can be generated by linear combinations of correlated traits. Corollary 1 combined with corollary 2-3 underlies the success of UBHDD since UBHDD only constrains the correlation between dependent variable (response) and independent variables (predictor) but not that among independent variables.

### Supplementary Note III

#### Phenotype space simulation

The basic parameters to conduct phenotype space simulation include the number of trait (*n*), the number of samples or population size (*m*=1000), the number of G-dimensions (*N*_1_=10, 20, 50, 100) and the number of NG-dimensions (*N*_2_=10,000), the number of G-dimensions per trait (*d*_1_=*N*_1_/2), the number of NG-dimensions per trait (*d*_2_=*N*_1_/2), broad-sense heritability (*H*^2^) or the variance of genetic component for standardized traits (*H*^2^ ~ U (0, 1)). A space can be expressed by a set of bases. When the set of bases are orthogonal, the dimensionality of the space is equal to the number of the set of bases. Assume *P*^G^ is independent of *P*^NG^. Then, the matrix composed of G-dimensions and NG-dimensions (*m* by (*N*_1_+*N*_2_)) is randomly generated by standard multivariate normal distribution (R package ‘MASS’). Then, the first *N*_1_ column vectors are set to be G-dimensions, the set of orthogonal bases of *P*^G^ subspace and the left *N*_2_ column vectors are set to be NG-dimensions, the set of orthogonal bases of *P*^NG^ subspace. For a focal trait (*T_i_*), the *P*^G^ component 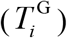 is formulated as

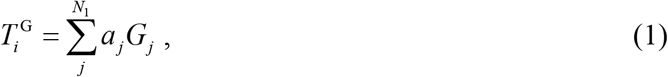

and the *P*^NG^ component 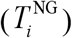 is formulated as

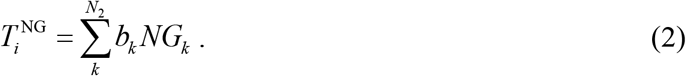

Then, 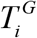 and 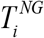 are standardized. Finally, the focal trait (*T_i_*) is generated by

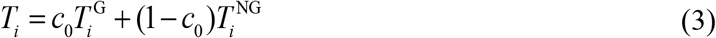

The coefficients *a_j_* and *b_k_* are randomly sampled from normal distribution N (0, 1) and random *N*_1_-*d*_1_ coefficients of *a_j_* and random *N*_2_-*d*_2_ coefficients of *b_k_* are set to be zero. The *c*_0_ satisfies

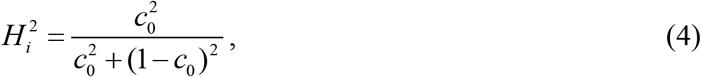

where 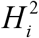 is the *H^2^* of *T_i_* assigned at the parameter setting. For each simulated *T_i_* defined by Eq. (3), we also define a set of correlated traits of *T_i_* as

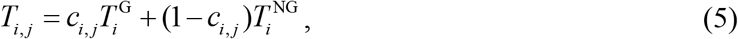

where *c_i,j_* is the coefficient controlling the j^th^ correlated trait (*T_i,j_*) of *T_i_* and satisfies

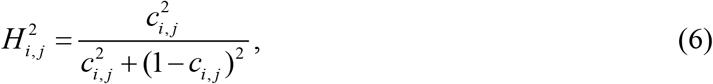

where 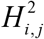 is the *H*^2^ corresponding to the j^th^ correlated trait (*T_i,j_*) of *T_i_* and satisfies

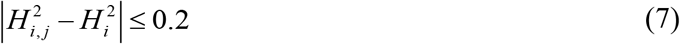

In an unstructured population, the number of correlated traits defined for each trait is the same (cluster size equal to 10 in the study). If we set different numbers of correlated traits to different traits, we can simulate a structured population (50 clusters of size 10, one cluster of size 200 and one cluster of size 300 in this study). For *N*_1_=10, 20, 50 and 100, we set *n*=1000 and *m*=1000; for *N*_1_=100, we also set *n*=2000 and *m*=1000 and *n*=2000 and *m*=2000 to observe apparent separation between *P*^G^ and *P*^NG^. Notably, *d*_1_ and *d*_2_ are set without loss of generality assuming that they are sufficiently small relative to *N*2.

**Table S1.**
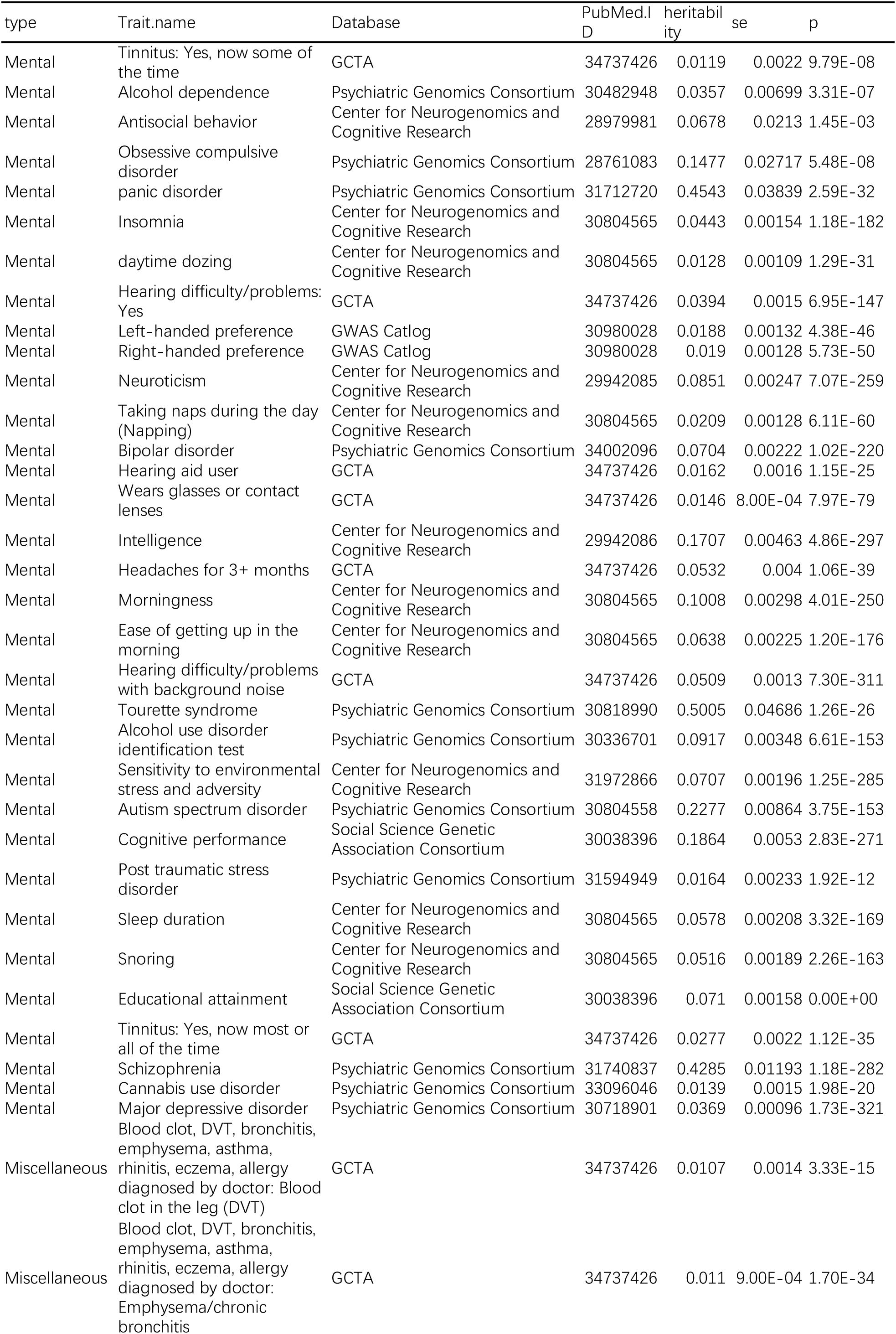
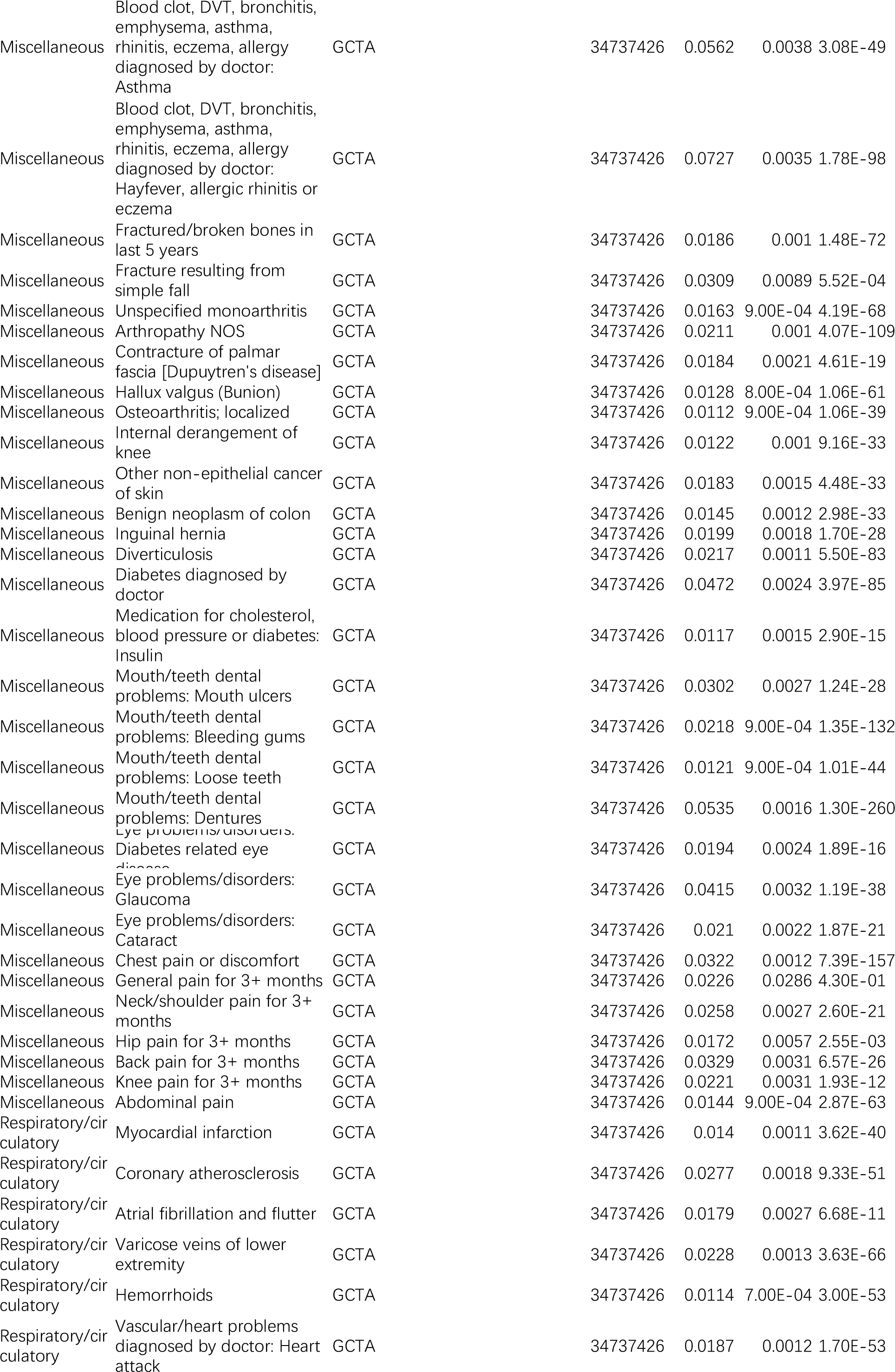
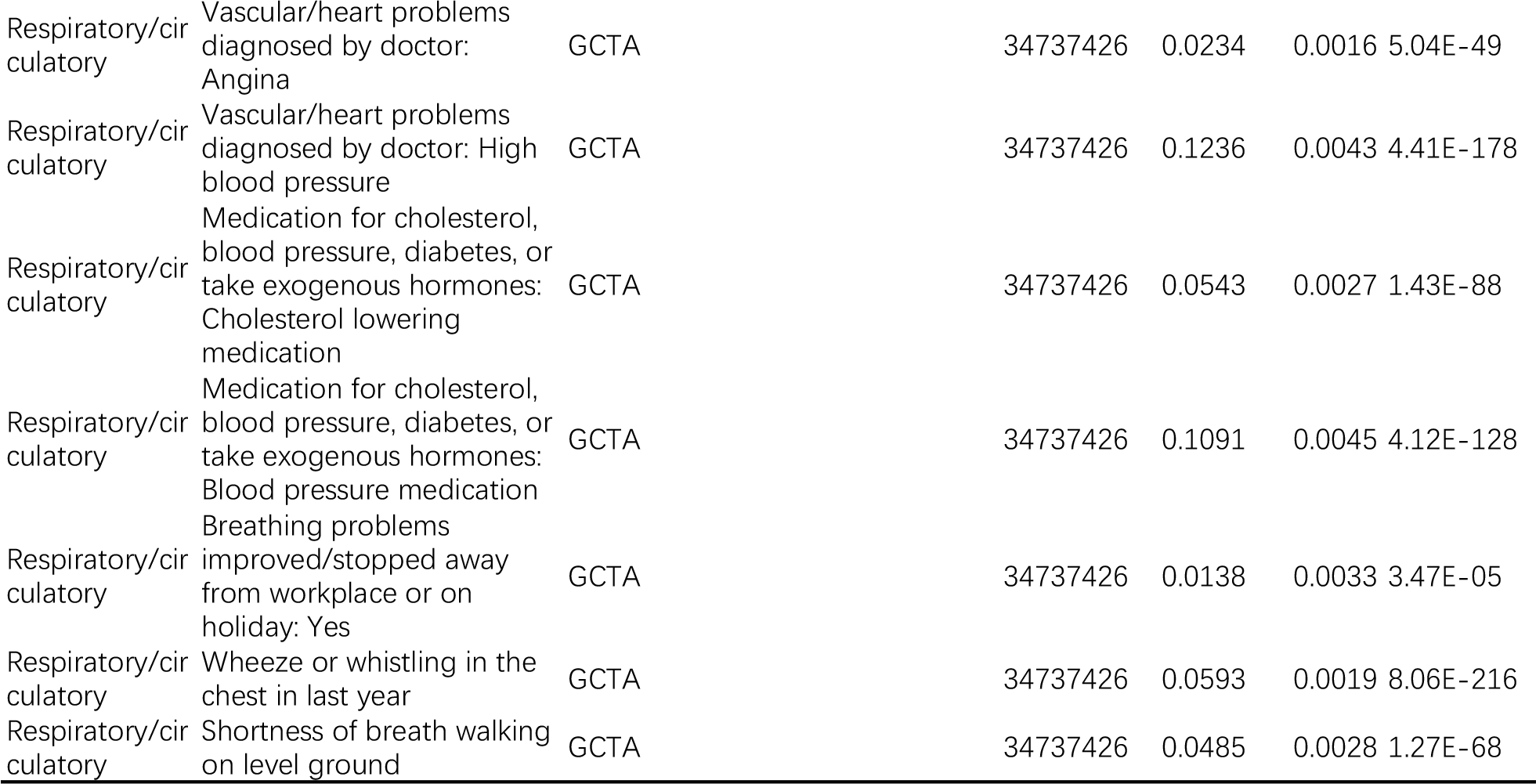
Curated mental traits/diseases

**Fig. S1.**
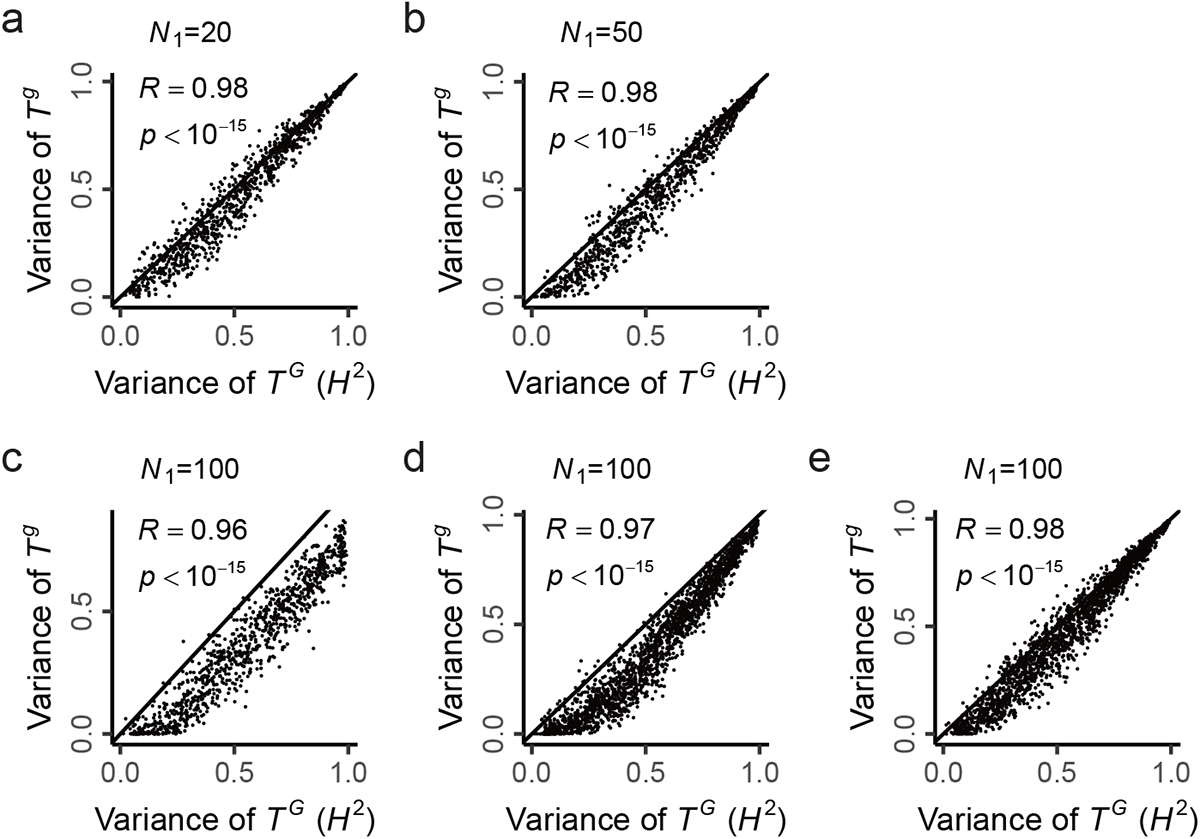
Decomposition of *P*^G^ and *P*^NG^ by UBHDD in simulated phenotype spaces. (a) The dimensionality of *P*^G^ subspace (*N*_1_) =20, polulation size (*m*) =1000 and the number of traits (*n*) =1000. (b) *N*_1_ =50, *m*=1000 and *n*=1000. (c) *N*_1_=100, *m*=1000 and *n* =1000. (d) *N*_1_ =100, *m*=1000 and *n* =2000. (e) *N*_1_ =100, *m*=2000 and *n* =2000. (c) shows a poor perfomance but can be improved with increasing number of traits (d) and further achieves higher performance with the increasing of population size (e).

**Fig. S2.**
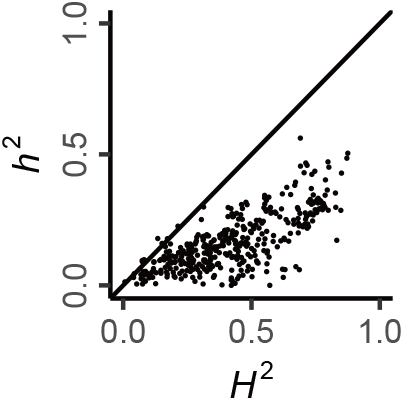
Broad-sense heritability (*H*^2^) and narrow-sense heritability (*h*^2^) of 405 yeast traits.

**Fig. S3.**
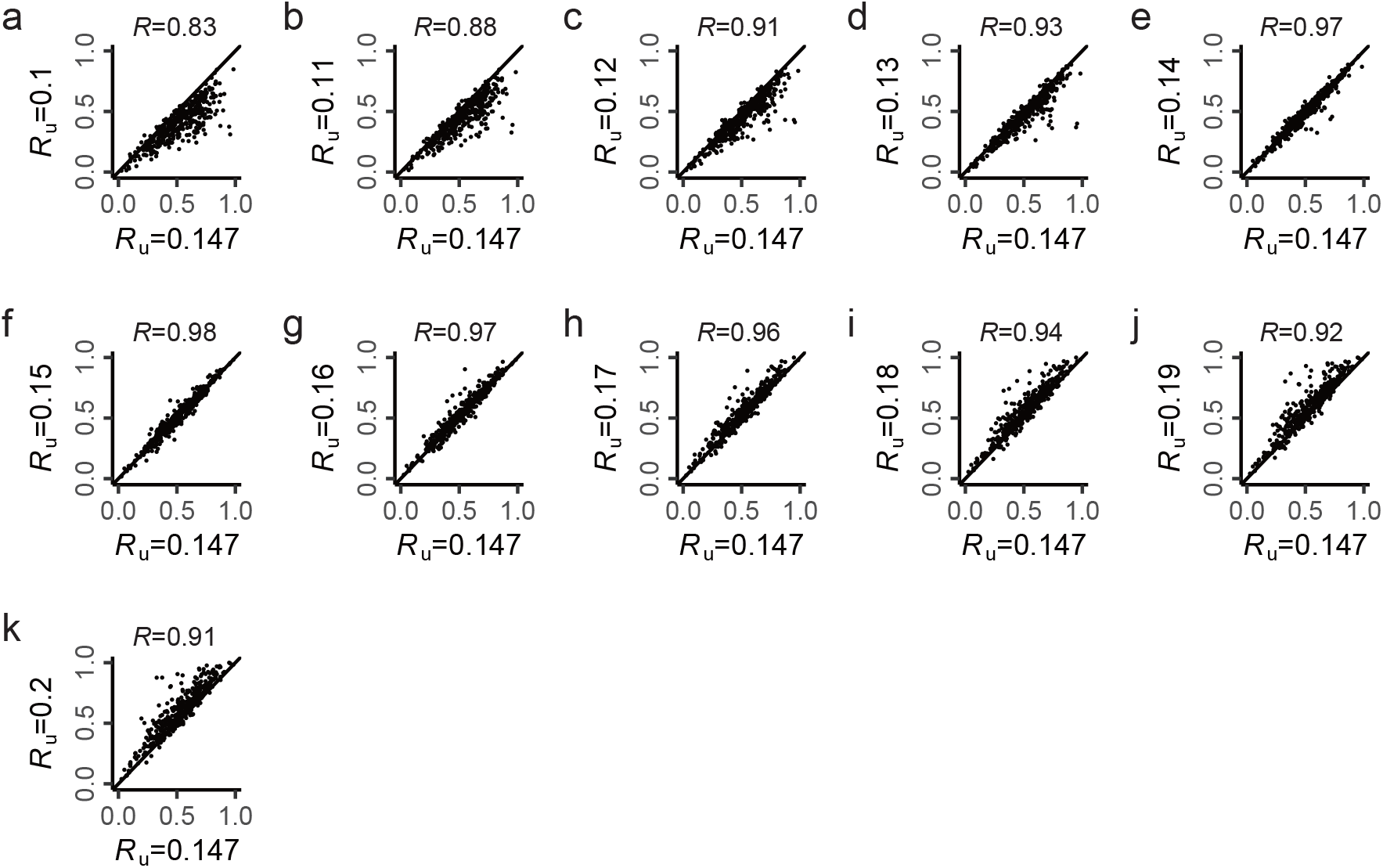
Robust estimation of *T^g^* by UBHDD under different uncorrelation thresholds (*R*_u_) in yeast. The threshold 0.147 corresponds to *p*=0.01 with Bonferroni correction in yeast seg-population. The estimated genetic variance is robust to the threshold used to conduct UBHDD.

**Fig. S4.**
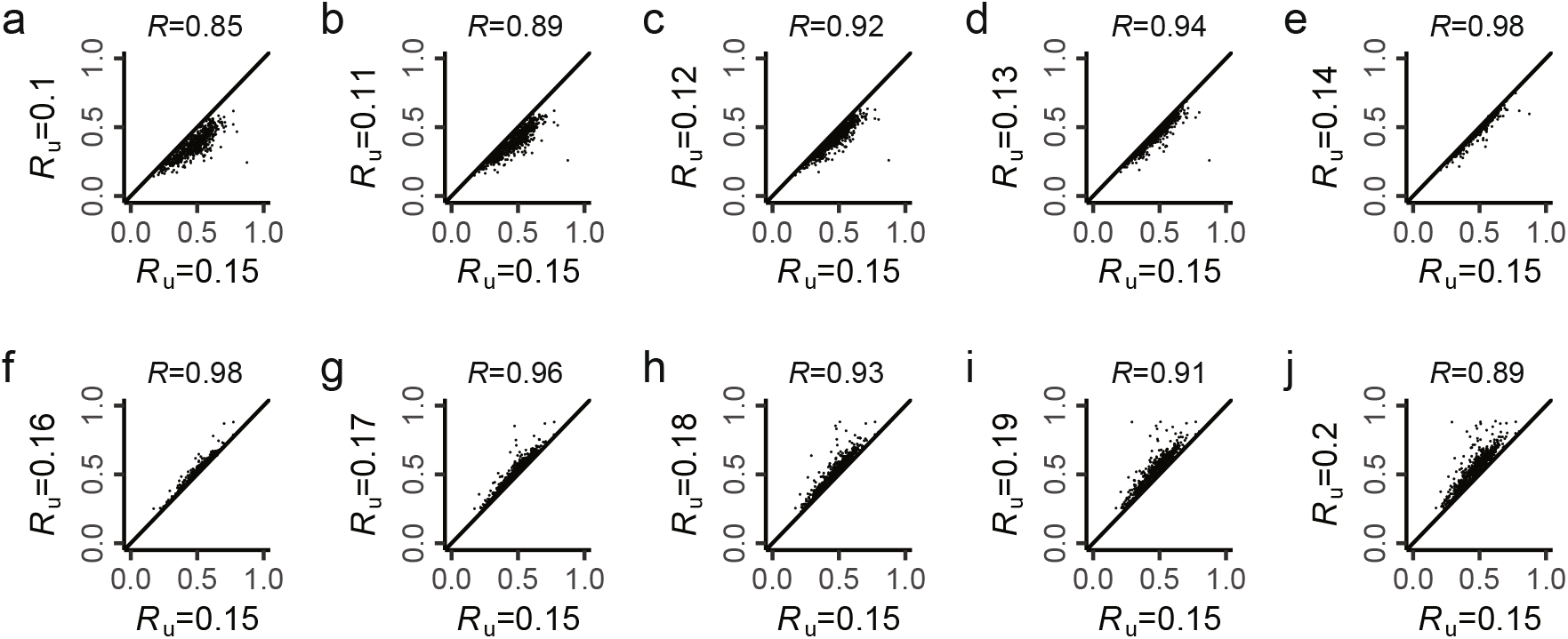
Robust estimation of *T^g^* by UBHDD under different uncorrelation thresholds (*R*_u_) in human brain. The threshold 0.15 is used in human brain phenotype space. The estimated genetic variance is robust to the threshold used to conduct UBHDD.

**Fig. S5.**
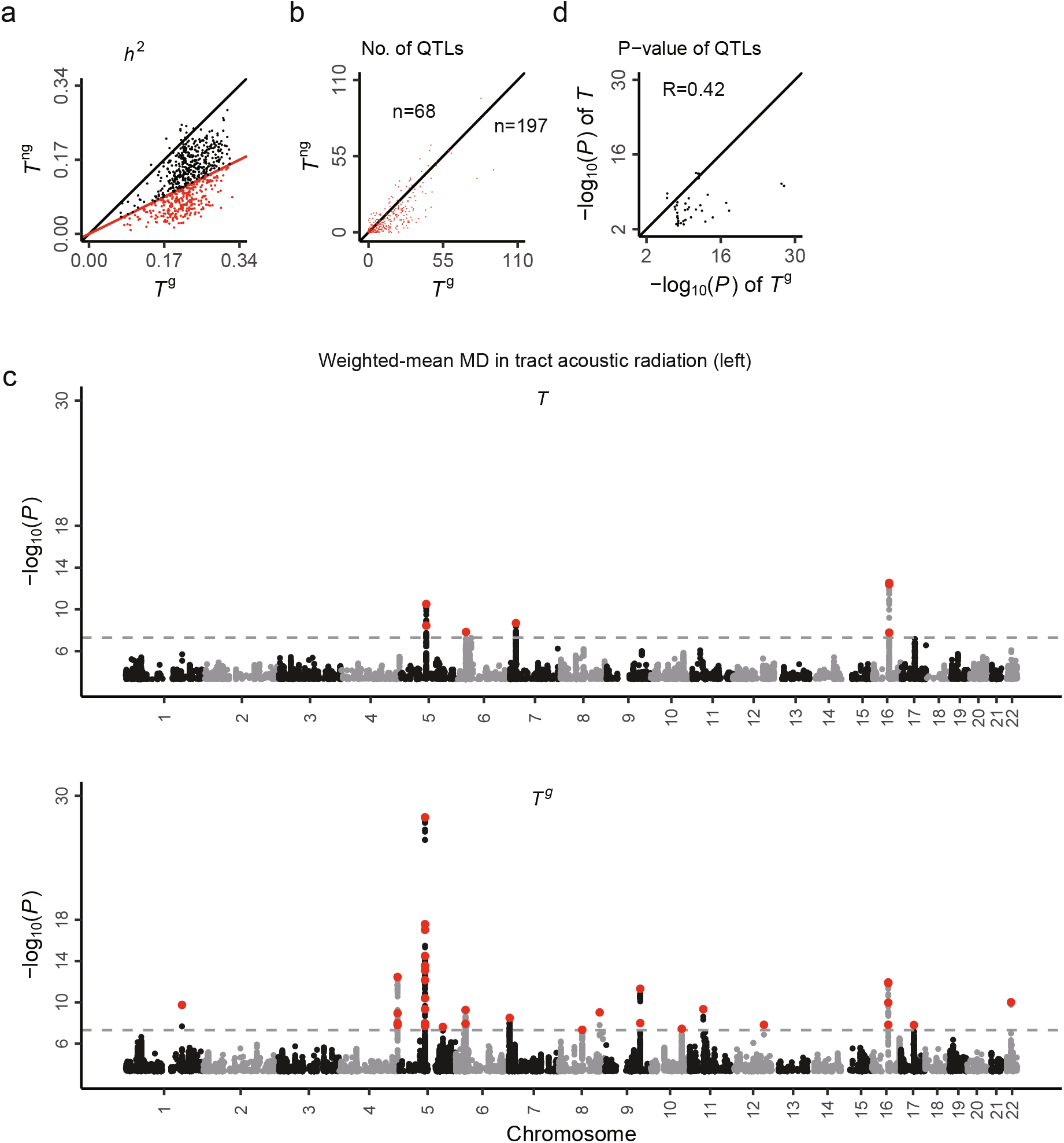
More QTLs found in *T*^g^ than in *T* for dMRI traits with strong enrichment of additive variance in *T*^g^. (a) Narrow-sense heritability (*h*^2^) is estimated by GCTA for 675 brain dMRI traits (R package ‘apcluster’). The *h*^2^ of *T*^g^ is generally larger than that of *T*^ng^. Red color shows the traits with at least two-fold enrichment. (b) The *T*^g^ generally has larger number of QTLs than *T* for traits with at least two-fold enrichments in (a). (c) The Manhattan plots of *T* and *T*^g^ are shown for an exemplar trait, weighted-mean MD in tract acoustic radiation (left). For the original trait *T*, 7 QTLs are mapped across 4 chromosomes but 35 QTLs across 13 chromosomes are mapped for the genetic component *T*^g^ estimated by UBHDD. The dotted line shows the threshold *p*= 5×10. (d) These extra QTLs in *T*^g^ often show strong but statistically insignificant signal in original trait *T*.

**Fig. S6.**
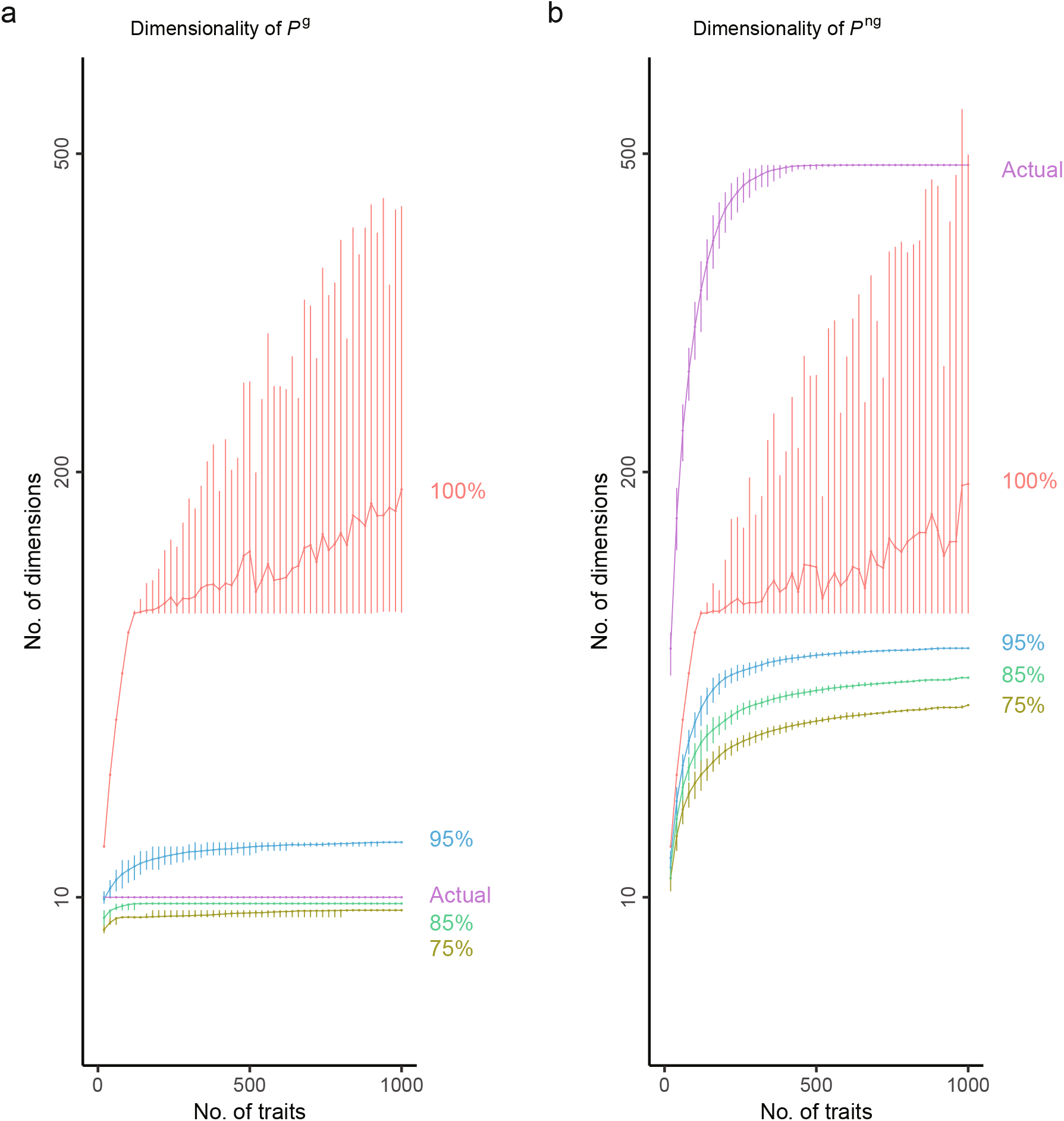
Dimensionality estimation of *P*^g^ and *P*^ng^ subspaces by PCA. We conduct PCA in *P*^g^ or *P*^ng^ and define the top PCs with 85% of variance explained as PC dimensions. Different cutoffs (75%, 85%, 95% and 100%) are compared. The error bars shown in lines represent 95% quantile of 100 sampling repeats. Middle lines represents the mean value. Notably, the number of PC dimensions in *P*^ng^ is always underestimated because PCA tends to merge independent dimensions in a population of small same size, especially when the dimensionality of *P*^NG^ subspace is larger than the rank of *P*^ng^ matrix. In the contrary, the PC dimensionality of *P*^g^ well approximates the actual dimensionality of the subsapce at the 85% cutoff. When larger cutoffs (95%, 100%) are chosen, the PC dimensions of *P*^g^ subspace will be overestimated. The overestimation happends because weak noise of modelling is falsely taken as dimensions. To facilitate comparison, the actual dimensionality in *P*^G^ or *P*^NG^ subspaces are plotted the same with Fig. 1f. The seemingly aberrant error bar at the 100% cutoff is also contributed by the PCA method (R function princomp return different number of PCs with 100% variance explained when traits are reordered.).

**Fig. S7.**
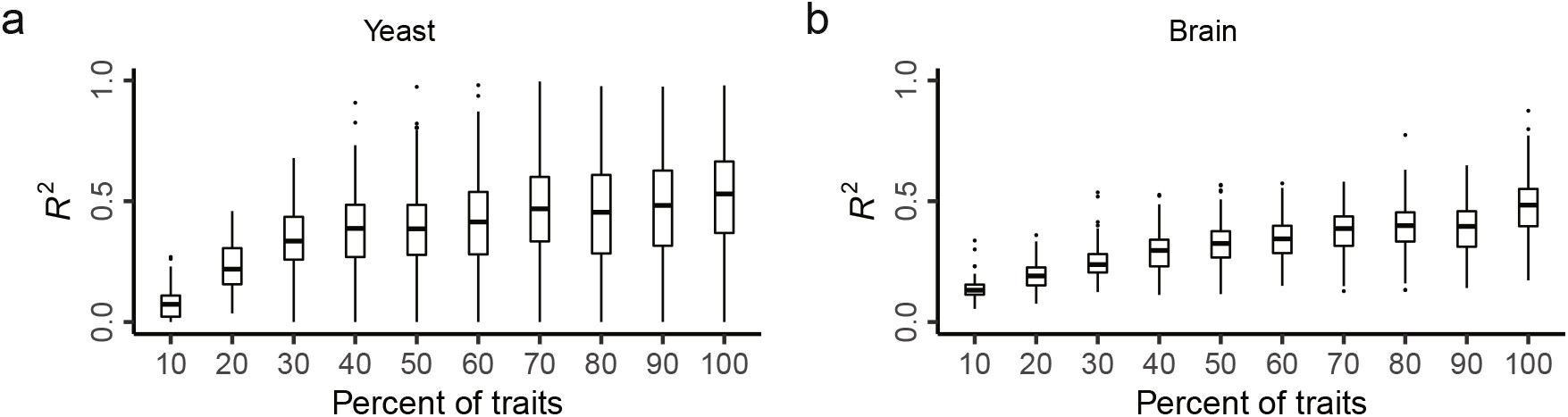
Evaluation for the number of traits saturated with G-dimensions in phenotype spaces. The accurate separation of genetic and non-genetic subspaces not only depends on rational uncorrelation threshold but also enough trait sampling. We conduct the same learning process for different proportions of trait subsets from 10% to 100%, say, a down-sampling strategy. Then, the distribution of UBHDD performance (*R*^2^, the variance of genetic component estimated by UBHDD) is compared among these trait subsets. (a) shows the distributions of yeast. (b) shows the distributions of human brain.

**Fig. S8.**
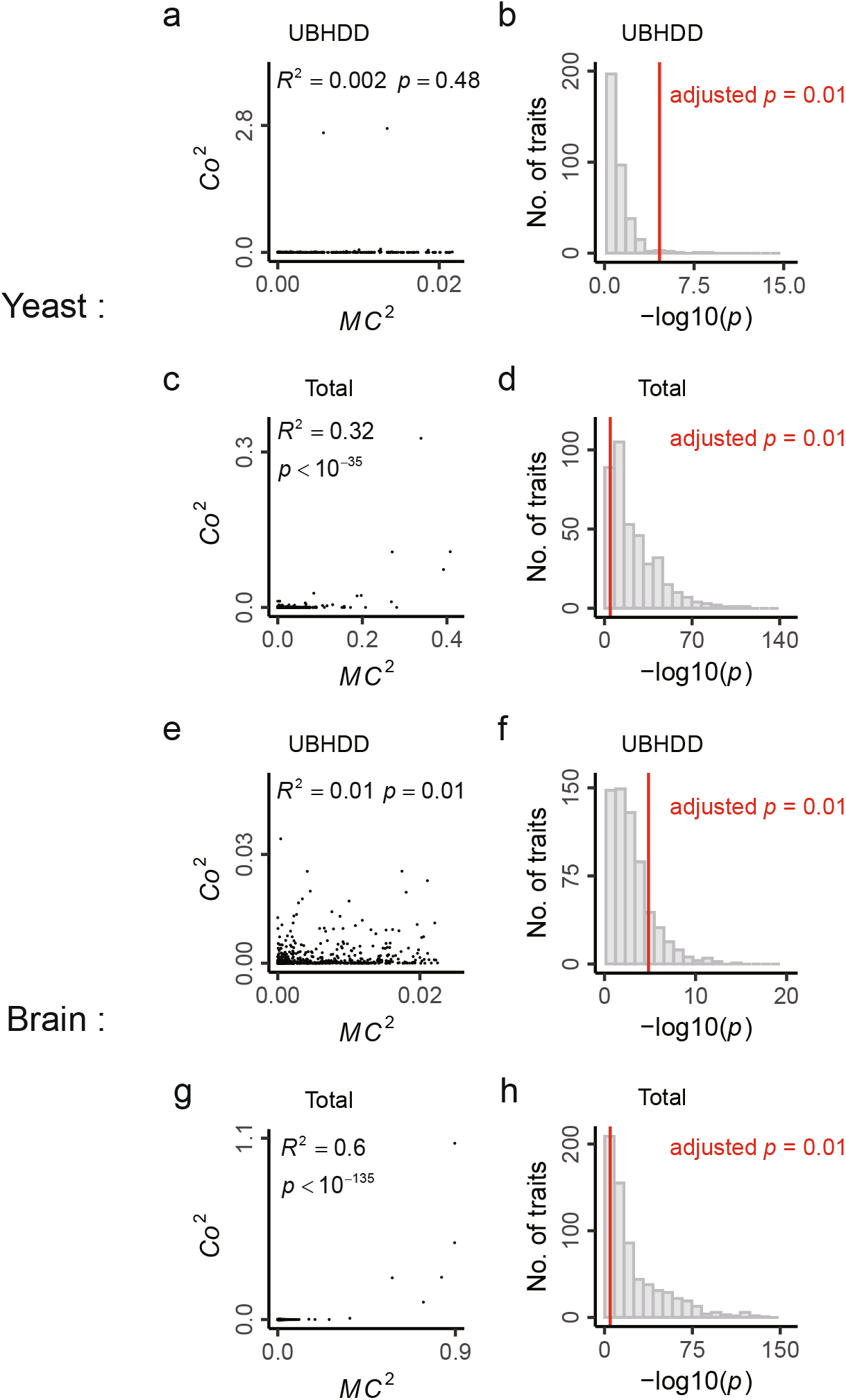
Criterion for uncorrelation threshold (*R*_u_). For a focal trait (*T_i_*), we can learn a linear function buit on its uncorrelated traits 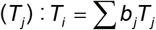. To judge the uncorrelation threshold (*R*_u_), we provide a statistical test as follows. First, we calculate the square of marginal correlation (*MC*^2^) between the focal trait and each of its uncorrelated traits. Then, we calculate the square of coefficients (*Co*^2^) for each uncorrelated trait in the learned linear function, say, 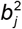. An optimal threshold is determinded if the *R*^2^ between *MC*^2^ and *Co*^2^ is insignificant, meanwhile, taking the number of uncorrelated traits available into account. (a-b) are results of UBHDD model. (a) shows the result under the *R*_u_ used in this study for an example trait in yeast. (b) shows the results for all of the 405 traits in yeast. As a contrast, we also learned linear functions based on total traits (Total model) for each of the 405 traits in yeast. (c-d) are results of Total model. (c) shows the same example trait in yeast based on Total model. (d) shows the results for all of the 405 traits in yeast based on Total model. Similarly, the results in human brain are shown for UBHDD model (e-f) and Total model (g-h). The red line denotes the adjusted *p*=0.01 to the number of traits.

